# Social evolution of shared biofilm matrix components

**DOI:** 10.1101/2021.12.16.472970

**Authors:** Jung-Shen B. Tai, Saikat Mukherjee, Thomas Nero, Rich Olson, Jeffrey Tithof, Carey D. Nadell, Jing Yan

**Affiliations:** Department of Molecular, Cellular and Developmental Biology, Yale University, New Haven, CT 06511, USA; Department of Mechanical Engineering, University of Minnesota, Minneapolis, MN 55455, USA; Department of Molecular Biology and Biochemistry, Wesleyan University, Middletown, CT 06459, USA; Department of Biological Sciences, Dartmouth College, Hanover, NH 03755, USA; Quantitative Biology Institute, Yale University, New Haven, CT 06511, USA

## Abstract

Biofilm formation is an important and ubiquitous mode of growth among bacteria. Central to the evolutionary advantage of biofilm formation is cell-cell and cell-surface adhesion achieved by a variety of factors, some of which are diffusible compounds that may operate as classical public goods – factors that are costly to produce but may benefit other cells. An outstanding question is how diffusible matrix production, in general, can be stable over evolutionary timescales. In this work, using *Vibrio cholerae* as a model, we show that shared diffusible biofilm matrix proteins are indeed susceptible to cheater exploitation, and that the evolutionary stability of producing these matrix components fundamentally depends on biofilm spatial structure, intrinsic sharing mechanisms of these components, and flow conditions in the environment. We further show that exploitation of diffusible adhesion proteins is localized within a well-defined spatial range around cell clusters that produce them. Based on this exploitation range and the spatial distribution of cell clusters, we construct a model of costly diffusible matrix production and relate these length scales to the relatedness coefficient in social evolution theory. Our results show that production of diffusible biofilm matrix components is evolutionarily stable under conditions consistent with natural biofilm habitats and host environments. We expect the mechanisms revealed in this study to be relevant to other secreted factors that operate as cooperative public goods in bacterial communities, and the concept of exploitation range and the associated analysis tools to be generally applicable.

## Introduction

Biofilms are an ancient and ubiquitous form of bacterial life in which cells form surface-attached groups embedded in an extracellular matrix^1^. Biofilm-dwelling bacteria gain survival advantages from the matrix including resistance to chemical and biological threats such as antibiotic exposure, host immune system defenses, and invasion and predation by other species^2–6^. Biofilm formation also contributes to the pathogenicity of many infectious bacterial species, and thus makes an outsized impact on human health^7–10^. The evolutionary advantage of biofilm formation is often tied to cell-cell and cell-surface adhesion, through which biofilm-forming species gain strength in numbers, resistance to physical disturbance, and physical association with favorable surfaces^11,12^. Indeed, in the native habitats of many biofilm-formers, nutrient distribution is heterogeneous and often localized around solid substrates^13–15^. Among the components of the extracellular matrix, exopolysaccharides and accessory proteins have been suggested to play dominant roles in biofilm adhesion to both biotic and abiotic surfaces in different species^3^.

In order to function properly, the adhesive molecules associated with biofilm matrix must be secreted outside of cells. This raises an important ecological question: *is the production of adhesive molecules exploitable, and if so, what conditions allow for their evolutionary stability?* In many contexts, secreted beneficial substances that diffuse away from producing cells can be scavenged by exploitative mutants (often termed cheaters), leading to a public goods dilemma^16,17^ in which cooperative behaviors partially or completely break down because producing cells, which pay a metabolic cost, are outcompeted by exploitative mutants^17–20^. The diffusive nature of secreted public goods necessarily leads to a characteristic length scale over which they can be shared, or exploited^21–24^. Previous literature has introduced the idea that secretion of public goods can be stabilized against exploitation if this length scale is smaller than or similar to the scale over which clonemates are clustered together, allowing cooperative cells to preferentially help each other relative to the population as a whole^25–29^. This principle is directly analogous to the classical statement of social evolution theory that cooperation can be favored by natural selection when the benefits of receiving cooperative help, weighted by the relatedness coefficient among givers and receivers of cooperation, exceed the costs of cooperation^30^.

Several studies have explored the competitive dynamics of partially shared matrix components in colonies on agar or pellicles at the air-liquid interface^31–35^, which can serve as excellent experimental platforms; but the continuous film of bacteria formed in these cases does not always correspond to scenarios in nature where isolated cell clusters are separated from each other in space. In many cases, biofilm-forming bacteria form cell clusters due to constraints on movement or mother-daughter cell adhesion, and these clusters may be sparsely or densely distributed in space depending on environmental conditions^36^. The physical size of these clonal clusters and the distance between them, therefore, represent two independent length scales. In principle, these two length scales should be compared with the diffusive length scale of the public goods to determine the physical conditions that stabilize cooperative product secretion against cheating.

The biofilm-producing pathogen and marine microbe *Vibrio cholerae* offers an excellent model for studies that may provide a close coupling of theory and quantitative empirical testing on the distance-dependent cooperative behavior. *V. cholerae* produces a suite of matrix components – including vibrio polysaccharide (VPS) and the proteins RbmA, RbmC, and Bap1 – that have been extensively characterized. VPS and RbmA are shared in a restricted fashion within secreting cell lineage groups and thus minimally susceptible to exploitation by non-producing cells^4,37–40^; this pattern has also been observed for major matrix components in *Pseudomonas aeruginosa* and *Bacillus subtilis*^33,34,41,42^. On the other hand, RbmC and Bap1, which are particularly important for adhesion of cell clusters to underlying surfaces, can diffuse far away from producing cells^37,43^. Here we establish that these two matrix proteins are exploitable by non-producing cells, creating an associated public goods dilemma. Competition assays between producers and non-producers of RbmC and Bap1 in static culture and with flow reveal the conditions under which populations of adhesion protein producers can resist invasion by cheating cells. We further show that exploitation of these matrix components takes place within a quantifiable spatial range, which depends critically on rates of diffusion and advection. As a result, sparse distribution of cell groups, and environmental flow that limits the spatial range of shared products, work jointly to limit exploitation of the diffusible matrix. This resolution links well to the ecology of *V. cholerae* biofilms in natural habitats and can be straightforwardly related to fundamental social evolution theory^30^.

## Results

### RbmC and Bap1 are exploitable public goods in *V. cholerae* biofilms

To study competition between diffusible matrix (RbmC and Bap1) producers and potential cheaters in *V. cholerae* biofilms, we used a strain background (*vpvC^W240R^*) locked in a high level of cyclic diguanylate that constitutively upregulates biofilm production machinery^44^. Using a constitutive biofilm-producing background allowed us to perform biofilm culturing experiments more readily while avoiding confounding factors from biofilm gene regulation, but our results can be obtained with a wild-type *V. cholerae* background as well (Fig. S1). We generated an isogenic mutant with double deletions of *rbmC* and *bap1* as the cheating strain, because of their functional redundancy in conferring surface adhesion^37,43,45^. The producer and cheater each expressed a different fluorescent protein to distinguish them from each other (Fig. S2). We first examined whether there is a fitness cost in producing RbmC and Bap1 by comparing producer and cheater growth rates in monoculture. As shown in Figure 1a, the growth rate of the producer is significantly lower than that of the cheater, demonstrating a substantial cost of producing these matrix components.

**Fig. 1.**
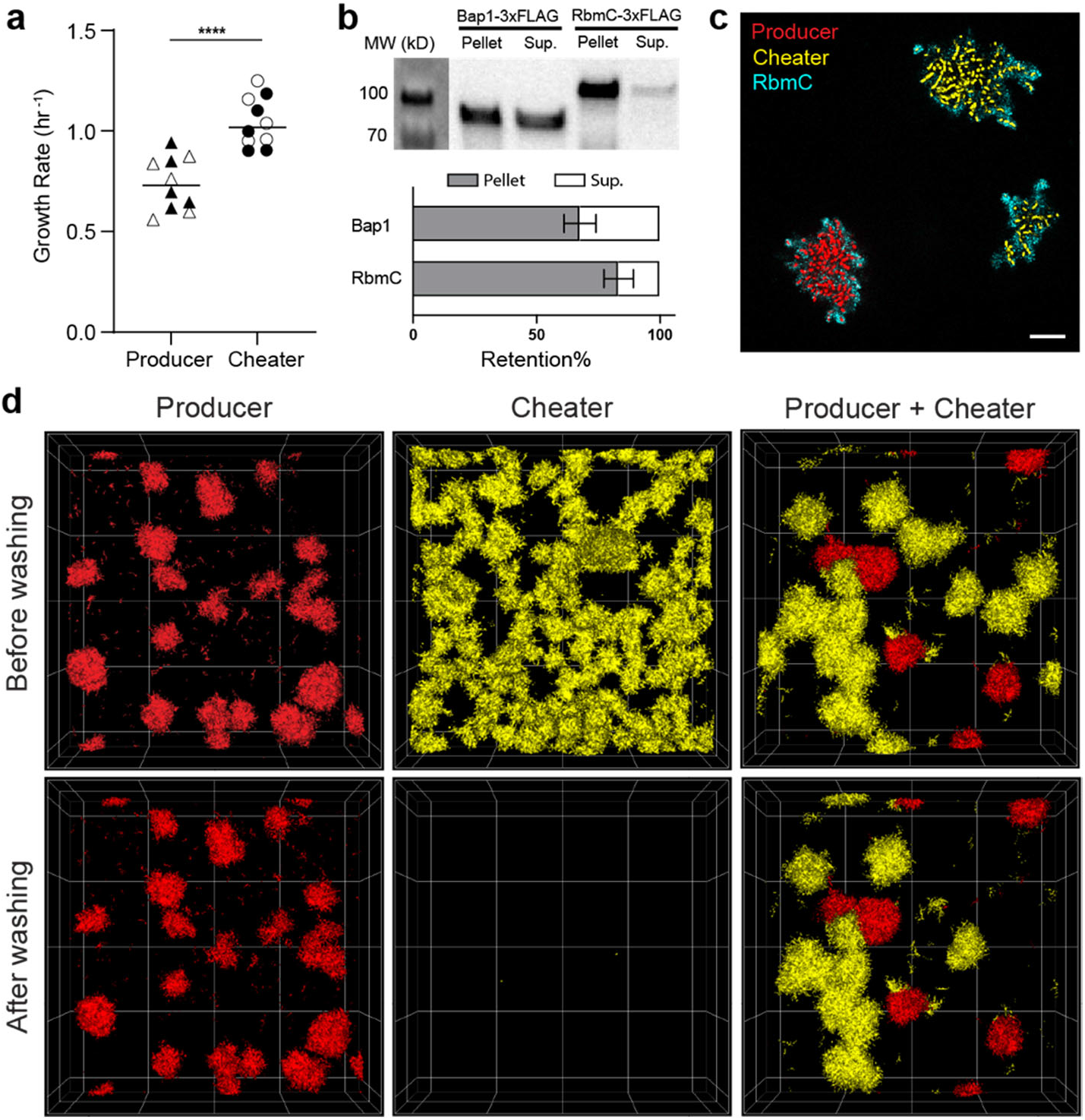
RbmC and Bap1 are exploitable public goods in *V. cholerae* biofilms. **a**, Growth rates of the producer strain and the cheater strain (Δ*bap1*Δ*rbmC*) from N = 10 independent replicates (****P < 0.0001, Mann-Whitney). Strains expressing mScarlet-I and mNeonGreen are shown by filled and empty markers, respectively. **b**, *Top*: Western blot showing Bap1 and RbmC in the biofilm (Pellet) and in the cell-free supernatant (Sup.). *Bottom*: quantification of the fraction of protein retained in biofilm (N = 3). **c**, A representative confocal image 7 μm above the substratum of a co-culture of producer (red) expressing 3×FLAG-tagged RbmC and cheater (yellow), stained with Cy3-conjugated anti-FLAG antibodies (cyan) to show the spatial distribution of RbmC. Scale bar: 10 μm. **d**, Mono-culture of producer (red), mono-culture of cheater (yellow), and co-culture of producer and cheater before and after washing. The producer constitutively expresses mScartlet-I and the cheater constitutively expresses mNeonGreen. Image size: 222 × 222 × 60 μm.

To document the degree of retention versus diffusive sharing of RbmC and Bap1, we performed a quantitative Western assay and found that 83% of RbmC and 68% of Bap1 were retained within producer biofilms, with the remaining protein detected in the surrounding liquid medium (Fig. 1b). Recapitulating prior literature, this result confirms that a substantial fraction of RbmC and Bap1 molecules can leave the cell clusters that produce them^37,43^. Next, to confirm that the molecules diffusing into the medium can be exploited by the cheaters, we tagged the C-terminus of RbmC or Bap1 with a 3×FLAG epitope and followed their spatial distributions in co-cultured biofilms using *in situ* immunostaining. The staining results show that significant amounts of RbmC and Bap1 are relocated from producer to cheater biofilm clusters (Fig. 1c and Fig. S3).

To determine whether exogenously acquired Bap1 and RbmC are functional in adhering cheater cell clusters to the substratum, we grew producer and cheater biofilms separately and in co-culture in 96-well plates. After growing biofilms statically for 16 h, we compared the biofilm biomass remaining on the glass substratum after a washing step that removes any weakly-adherent cell clusters (see Methods) (Fig. 1d). We found that in monocultures, almost all producer cell clusters remained surface-bound after washing, whereas nearly zero biomass was found for cheater biofilms after the disturbance, consistent with the absence of surface adhesion in the cheater strain. In contrast, in the co-culture of producer and cheater, some of the cheater cell groups were able to resist the strong fluid flow during the washing step. Coupling this observation with the immunostaining results, we inferred that adhesion can be partially restored in cheater biofilms by taking advantage of the adhesion proteins produced by the cohabitating producers. Overall, these results confirm that the shared matrix proteins RbmC and Bap1 in *V. cholerae* biofilms are cooperative products that draw a significant cost to produce and are susceptible to cheater exploitation.

### Exploitation range and community spatial structure jointly determine the outcome of competition between producers and cheaters

Because RbmC and Bap1 are costly to produce but exploitable by cheating strains, it is possible that non-producing cells could outcompete producing cells in co-culture. To test if this is true, we first competed the producer against cheater in a static environment. The initial biomass of both strains was quantified by the inoculation number densities (*σ*_0,p_ and *σ*_0,c_ for producer and cheater, respectively; *σ*_0_ ≡ *σ*_0,p_ + *σ*_0,c_ is the total inoculation number density). After 16h of growth, the total biomass in three-dimensional (3D) space was quantified both before and after washing (Methods; Fig. S4). The competition assay was initiated at different initial frequencies of the producer *f*_0,p_ ≡ *σ*_0,p_/(*σ*_0,p_ + *σ*_0,c_) for a range of total inoculation number density *σ*_0_. Representative images and quantitative outcomes of experiments are shown in Fig. 2a-d, where Δ*f*_1,p_ ≡ *f*_1,p_ - *f*_0,p_ and Δ*f*_2,p_ ≡ *f*_2,p_ - *f*_0,p_ are the frequency changes of the producer compared to *f*_0,p_ before and after washing, respectively.

**Fig. 2.**
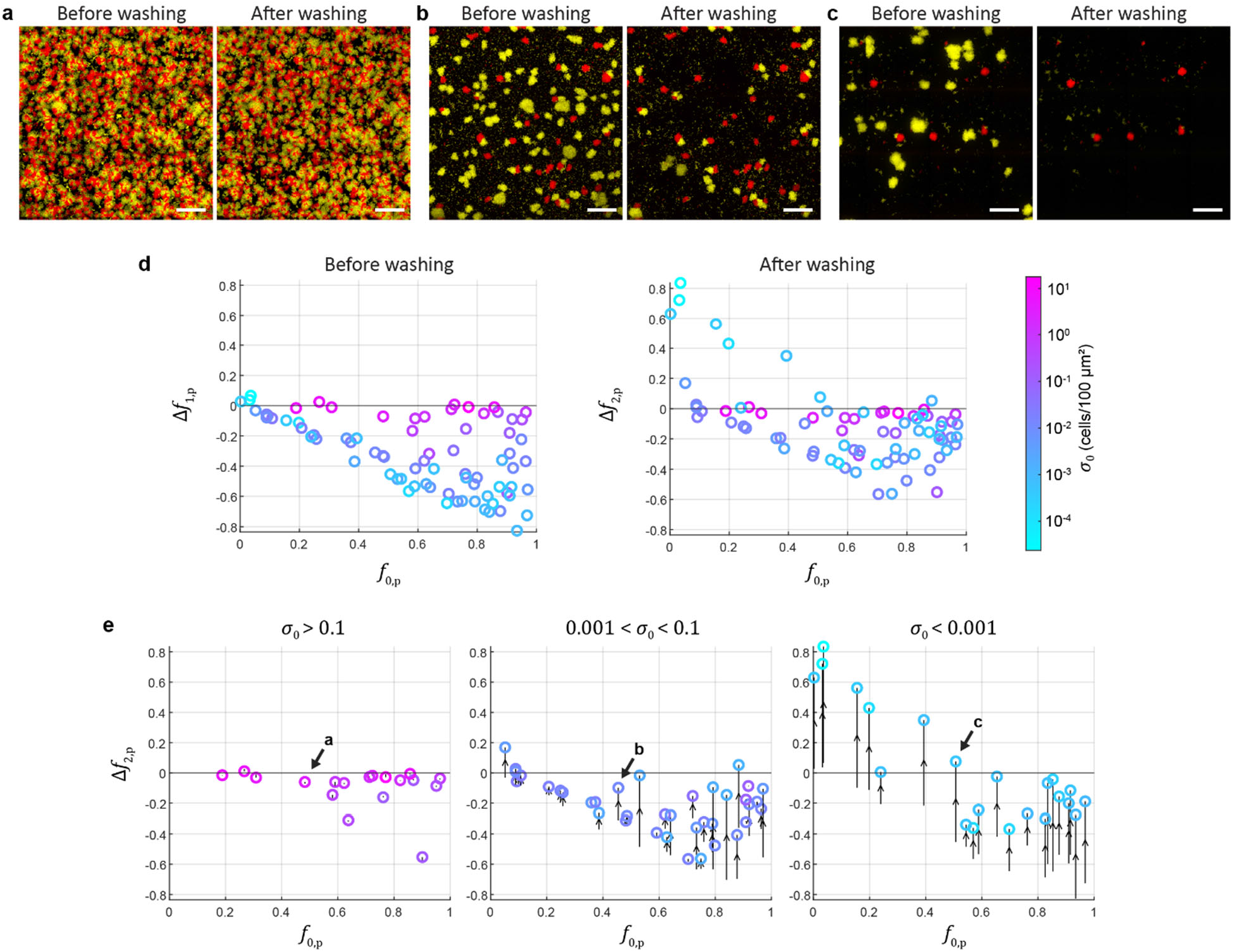
Competition between adhesion protein producer and cheater strains in a static environment. **a-c**, Producer (red) and cheater (yellow) biofilms grown in a static environment before and after washing with different inoculation number densities *σ*_0_ corresponding to data labeled in (e). Scale bars: 100 μm. **d**, Adhesion protein producers in competition with cheaters before and after washing. **e**, Adhesion protein producers in competition with cheaters in different *σ*_0_ ranges (color-coded) in a static environment. Each data marker denotes the frequency change in producer after washing (Δ*f*_2,p_), and the corresponding frequency change before washing (Δ*f*_1,p_) is connected to it by an arrow (Δ*f*_1,p_ → Δ*f*_2,p_).

We found that before washing, cheaters outcompete producers at all initial frequencies, and the fitness of the producer decreases as the inoculation density *σ*_0_ decreases (Fig. 2d). This is because in a static environment, both producers and cheaters compete for the same nutrient pool, and the cheater has a substantial advantage in growth rate (Fig. 1a); also, importantly, the benefit of shared adhesion protein production to biofilm cells does not manifest in a disturbance-free environment. The decrease in producer fitness with decreasing *σ*_0_ is due to finite resources in the static environment: At high *σ*_0_, both strains can only undergo a short period of growth before reaching the environment’s carrying capacity, limiting the total change in relative abundance of the two strains that can occur. A classic resource-limited competition model reproduced the competition results in a static environment (Fig. S5).

In the competition data after washing, the producer fitness is systematically increased compared to the competition before washing, suggesting that cheater biomass is preferentially removed compared to producer (Fig. 2d). Importantly, this pattern shifted with *σ*_0_ and corresponded to negative frequency-dependent selection for RbmC and Bap1 production at *σ*_0_ < 0.001: at low initial coverage, the producer is positively selected at low initial frequency. This is evident in Fig. 2e where the competition data are sorted based on inoculation number density *σ*_0_. We expect that at even lower inoculation number densities, the stable point will move to *f*_0,p_ = 1, i.e. the producer will always win regardless of the initial frequency; obtaining sufficient statistics at such low *σ*_0_ is however experimentally difficult due to limits on field-of-view acquisition with sufficient detail to make accurate measurements. We also note that the magnitude of Δ*f*_2,p_ - Δ*f*_1,p_, the fitness difference in producer before and after washing, increases with decreasing *σ*_0_ (Fig. 2e), hinting at the importance of the distance between producer and cheater clusters for the effect of shear disturbance on population dynamics.

To understand the mechanism underlying the sharing dynamics in the competition assay, we analyzed how exploitation of diffusible matrix adhesion proteins takes place in a spatial context. We observed at low *σ*_0_, cheater biomass is distributed predominantly around isolated producer biofilms after washing (Fig. 3a). Based on this observation, we hypothesized that there exists a finite spatial range in which cheater cell clusters are protected against fluid disturbance by RbmC and Bap1 secreted by the producer (Fig. 3b). We define the radius of this range as the exploitation radius *R*, which we extract by calculating the cross-correlation between the producer and cheater biomass distributions in the co-cultures after washing (Fig. 3c). Conceptually, the farther away a cheater cluster is from a producer cluster, the less likely it persists on the substrate after washing – this leads to a decaying likelihood of finding cheater biomass around producers with increasing distance. By assuming an exponentially decaying cross-correlation, we arrived at an exploitation radius *R* value of 49 ± 11 μm (Figs. 3c and S6; Methods). Note that this value is significantly larger than the physical size of the producer cell cluster, *r*_0_ (≈ 9 μm). This value agrees with the visual inspection of cheater distribution in Fig. 3a. Using this method, we also confirmed the partial redundancy of RbmC and Bap1 in providing adhesion between surfaces and cell clusters, as both single mutants give a similar exploitation radius (Fig. S7).

**Fig. 3.**
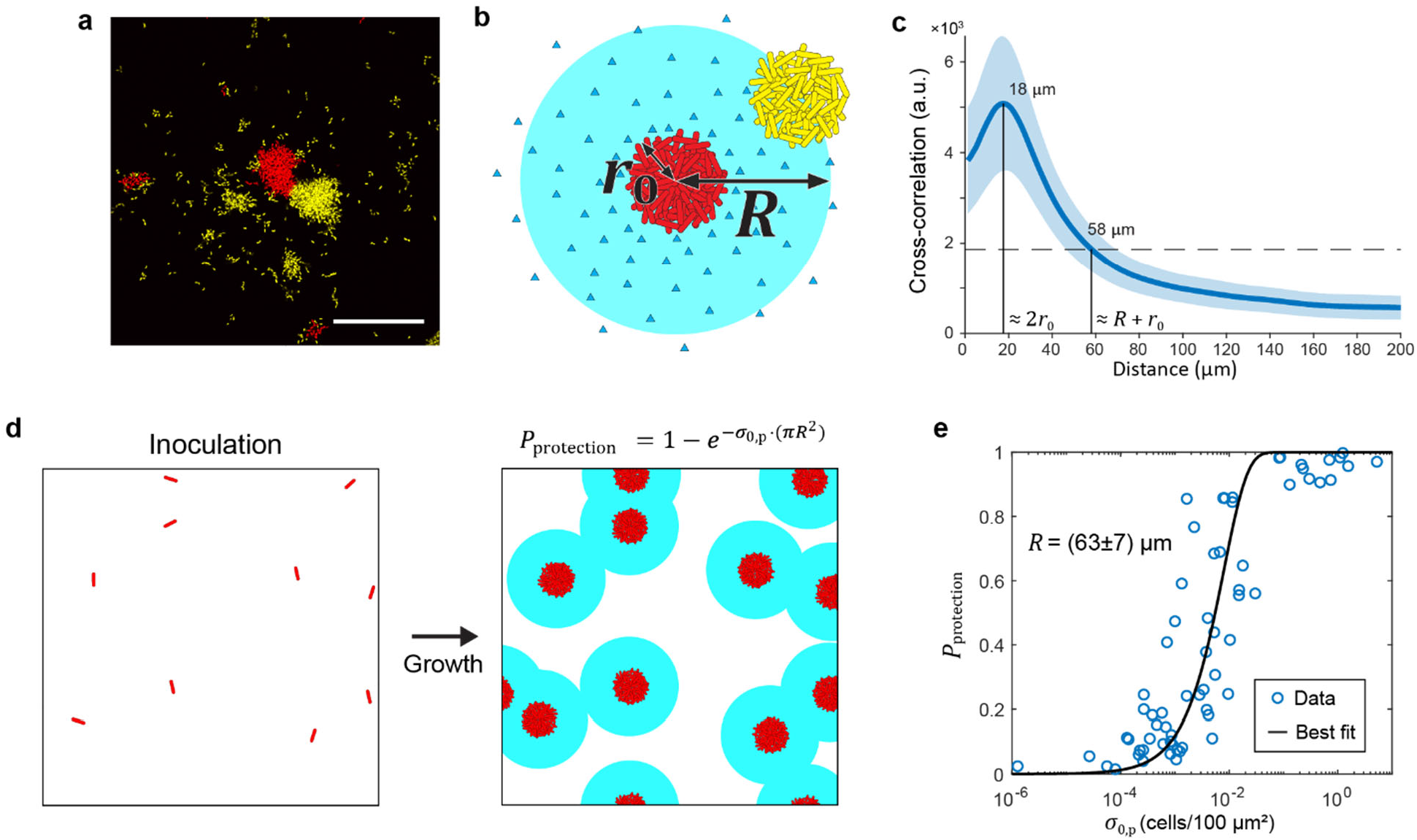
A spatial exploitation model quantitively explains competition dynamics. **a**, Representative confocal image of a co-culture of producer (red) and cheater (yellow). The maximum intensity projection image shows the distribution of cheater biomass around isolated producer clusters after washing. Scale bar: 50 μm. **b**, Schematic of exploitation radius *R* provided by diffusion of adhesion proteins from a producer cluster. *r*_0_ is the physical radius of the producer cluster. **c**, Exploitation radius determined as *R* = 49 ± 11 μm by analyzing the cross-correlation between the distribution of producer and cheater biomasses. Error represents the standard error from N = 4 independent replicates. The peak at 18 μm corresponds to 2*r*_0_. The horizontal dashed line indicating *e*^−1^ of the maximum cross-correlation intersects at 58 μm, corresponding to *R* + *r*_0_. **d**, Schematic of the spatial model quantifying the protection probability *P*_protection_ conferred by randomly distributed producer clusters. Each inoculated individual cell contributes to an area of protection around it and all producer clusters collectively yield *P*_protection_ = 1 − exp (-*σ*_0,p_ ⋅ *πR*^2^), where *σ*_0,p_ is the inoculation number density of the producer. **e**, Experimentally measured *P*_protection_ plotted vs. *σ*_0,p_. Fitting the data with the equation in (d) yields *R* = 63 ± 7 μm with 95% confidence bounds.

Next, to cross-check the value of exploitation radius *R* and to show how this concept can quantitatively explain the competition dynamics, we constructed a spatial model to interpret the dependence of competition outcome on inoculation number density and initial producer frequency (Methods). Briefly, we assume that each producer cell cluster originates from a randomly distributed founder cell, inoculated at a number density *σ*_0,p_ (Fig. 3d). As such, cheaters within a distance *R* from any producer cluster remain adherent after washing; otherwise, they are removed. In this setting, the probability *P*_protection_ of a cheater cluster falling into the exploitation radius of any producer cluster and being protected from flow disturbance is related to the total exploitable area provided collectively by all of the producers in the system. To calculate the latter, we treat each producer cell cluster as contributing a disk of area *πR*^2^ of protection (Fig. 3d). Note that these imaginary disks can overlap with each other, because there is no constraint on the distance between inoculated producer cells. The relation between *σ*_0,p_, *R* and *P*_protection_ can be derived from probability theory: *P*_protection_ = 1 − exp(−*σ*_0,p_ ⋅ *πR*^2^) (Fig. 3d; Methods). Importantly, *P*_protection_ can be easily extracted from the biomass data before and after washing by identifying *P*_protection_ as the portion of cheater biomass remaining adherent after washing (Fig. S4). Fitting *P*_protection_ vs. *σ*_0,p_ from experimental data with the theoretical relation yields *R* = 63 ± 7 μm, where the error corresponds to 95% confidence bounds (Fig. 3e). This value is slightly higher than the value extracted above (49 ± 11 μm), but the agreement between the two estimates – one from direct quantification of biomass distribution and the other derived from the spatial exploitation model, is remarkable. The two methods serve as a consistency test for each other and corroborate our hypothesis that exploitation of matrix proteins can take place beyond the physical size of the clonal cluster but only within a finite, well-defined length scale. A two-part model combining the spatial exploitation model with the structureless competition model can qualitatively reproduce all the experimental data in the 96-well plate (Fig. S5).

### Flow shifts the competition between producer and cheater by modulating the exploitation range of shared adhesion proteins

Thus far, we have studied the population dynamics of RbmC/Bap1-producing and cheating strains in static liquid environments, but in natural contexts of *V. cholerae* and many other microbes, including on marine detritus (so-called “marine snow”)^46^, biofilms grow under continuous fluid flow. To understand how a continuous flow may shift the competition between diffusible matrix producers and cheaters, we conducted competition assays in microfluidic channels with a flow rate of 1 μL/min, corresponding to an average flow speed of 694 μm/s (Methods). The flow speed falls into the range of the measured speed of sinking marine snow particles, the natural context of (non host-associated) *V. cholerae* and many other marine bacteria^47,48^.

The presence of flow dramatically changed the population dynamics relative to static co-cultures: competition was generally neutral at medium to high inoculation density (*σ*_0_ > 0.01), indicating that RbmC/Bap1-producer and cheater strains can coexist under this condition (Fig. 4, a-d). In contrast, at *σ*_0_ < 0.01, producers outcompeted cheaters regardless of initial population composition. Compared to a static environment, continuous flow generally enhanced the fitness of the RbmC/Bap1-producer against cheating strains (Fig. 2e). For example, in the range 0.01 < *σ*_0_ < 0.1 and *f*_0,p_ ≈ 0.5, an increase in the flow rate from 0 to 1 μL/min increased the fitness of the producer, an effect that saturates beyond 0.5 μL/min (Fig. 4e).

**Fig. 4.**
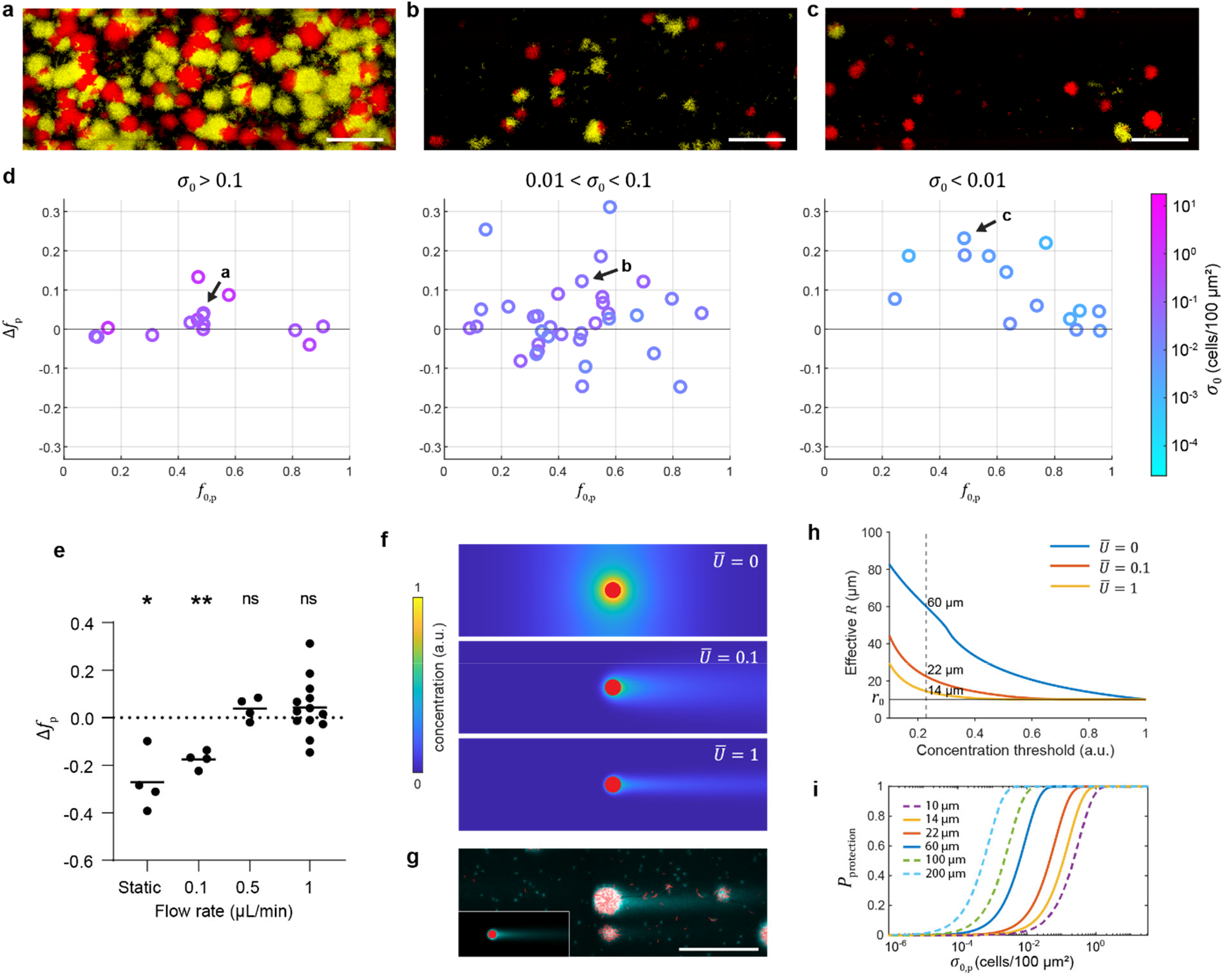
Flow shifts the competition between producer and cheater by modulating the exploitation range of the adhesion protein. **a-c**, Representative maximum intensity projection images of producer (red) and cheater (yellow) biofilms grown in a microfluidic channel with different *σ*_0_ corresponding to data labeled in (d). Scale bars: 100 μm. **d**, Producers in competition with cheaters at different *σ*_0_ in a flow environment (flow rate = 1 μL/min). **e**, Dependence of Δ*f*_p_ on flow rate. Data labeled static were obtained from the static competition assay in 96-well plate. Data were taken at 0.01 < *σ*_0_ < 0.1 and 0.4 < *f*_0,p_ < 0.6. One sample *t*-test was used to examine if Δ*f*_p_ is statistically different from 0 (*P < 0.05 and **P < 0.01; N = 4, 4, 4, and 13). **f**, Simulated depth-averaged adhesion protein concentration in a flow chamber at different average flow speeds 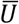. The concentration is normalized by the value at the edge of the producer cluster at 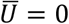. The red disk corresponds to a producer cluster of radius *r*_0_ = 10 μm. **g**, Distribution of adhesion protein Bap1 (cyan) around producer clusters (red) at the surface of the flow chamber shown by Cy3-conjugated anti-FLAG antibody staining. The corresponding simulation result is shown in the inset (*Lower left*). Scale bar: 100 μm. **h**, Effective *R* as a function of concentration threshold for protection at different flow rates in simulations. The horizontal line indicates the edge of the producer cluster at *r*_0_. A threshold concentration of 0.23 (vertical dashed line) yields *R* = 60, 22, and 14 μm for 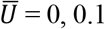, and 1, respectively. **i**, *P*_protection_ vs. *σ*_0,p_ calculated for different *R* values.

To understand mechanistically how fluid flow shifts the competition between RbmC/Bap1 producers and cheaters, we performed numerical modeling of the fluid flow and adhesion protein concentration around a producer cluster attached to the surface at different average flow speeds 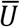 (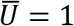 in simulations corresponds to 1 μL/min in experiments). The producer cluster is modeled as a surface-attached hemisphere and secretes matrix molecules at a constant rate from the biofilm-fluid interface (Methods). The simulation results show that, compared to the purely diffusive case, the adhesion protein concentration is drastically reduced around the producer cluster due to advection at a finite flow speed (Fig. 4f). Furthermore, the advective transport results in a comet-tail profile of the protein concentration at the glass surface along the wake of the fluid flow, which we confirmed in the immunostaining experiment performed under the same condition (Fig. 4g).

To explain the difference in fitness of the producers in the static and flow environments, we again applied the concept of exploitation range. Assuming a threshold concentration above which cheater biofilms receive sufficient adhesion proteins to resist flow, the corresponding effective *R* around producer cell clusters can be derived based on the simulated concentration profiles of RbmC/Bap1 at different flow speeds (Fig. 4f). As 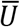 increases, the effective *R* reduces sharply: for example, setting an arbitrary threshold, as shown in Fig. 4h, yields *R* = 60, 22, and 14 μm for 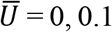, and 1, respectively. The ensuing *P*_protection_, which is dependent on *R*, decreases dramatically with decreasing *R* as shown in Fig. 4i. Our simulation results thus show that flow shrinks the effective exploitation radius conferred by an individual producer cluster^27^, which in turn reduces *P*_protection_ and positively shifts the competitive dynamics toward a regime that increasingly favors RbmC/Bap1-producing cells. The dependence of *P*_protection_ on flow therefore qualitatively accounts for the dependence of fitness on flow (Fig. 4e). Combined with the distance between producer cell clusters driven by sparse single cells colonizing the surface, flow limits the spatial range of sharable matrix components and averts exploitation of sharable matrix adhesion proteins.

## Discussion

Cell-cell attachment and cell-surface adhesion are often critical to the evolutionary advantage of biofilm formation. The extracellular matrix plays an important role in controlling biofilm organization and adhesion^37–39,45,49–54^, and understanding the evolutionary dynamics of its production remains an active area of work in this research domain. The biofilm matrix often contains components that vary in their diffusion properties and potential for exploitation by cells that do not produce them. In this study, we explore the population dynamics of diffusible matrix protein production in *V. cholerae* biofilms. We carried out competition assays under static and flow conditions, both of which are frequently encountered by *V. cholerae* in native habitats and in hosts^47,48,55^. We found that the matrix components RbmC and Bap1, as diffusible public goods, can be exploited and confer resistance to physical disturbance to clusters of non-producing cells within a finite spatial range around producer cell clusters. This exploitation range depends on the diffusion-advection condition of the environment, where a continuous flow effectively shrinks the exploitation range and consequently suppresses cheater exploitation. In their marine and fresh water reservoirs, *V. cholerae* cells often attach to floating food particles and experience shear flows ^47,48^. During infection, *V. cholerae* biofilms are primarily localized on the tips of intestinal villi and therefore experience peristaltic flow in the gut^55^. The flow speed and the associated shear stress used in this study fall within the range measured in these environments^56^. The population densities of *V. cholerae* cells in natural ecosystems have been measured to range from 10^2^ to 10^5^ cells per liter^57^, corresponding to an average distance between cells on the order of millimeters in 3D space. Thus, our results suggest that the combination of flow and low cell density drive the competition dynamics in favor of RbmC and Bap1 producers and lead to the evolutionary stability of these diffusible matrix proteins in *V. cholerae*.

It is interesting to contrast our results with those for the other major matrix protein in *V. cholerae* biofilms, RbmA, which (together with VPS) is responsible for high density cell-cell packing within biofilm cell clusters. RbmA is secreted and shared in a limited fashion within cell lineage groups producing it^4^, conferring protection from exploitation with little dependence on the distance between clusters of producing versus non-producing cells^4^. We envision that the different sharing dynamics of RbmA, RbmC, and Bap1 are constrained by their function as components of the biofilm matrix. RbmA, which holds mother-daughter cell lineages together, must be sequestered and stay in close proximity to producing cells to perform this function. RbmC and Bap1, on the other hand, must travel a distance away from producing cells to confer adhesion between cell groups and underlying surfaces, making them inherently sharable and exploitable; the exact biochemical and biophysical mechanisms underlying this difference await future research. As a consequence, different matrix components – even those produced by cells of one species within the same biofilm – can have different environmental constraints and population structures as conditions for their evolutionary stability.

Our results have a straight-forward correspondence with social evolution theory. Hamilton’s rule provides the canonical condition under which cooperation is favored by selection: the fitness benefit of receiving cooperative help, weighted by the relatedness coefficient that quantifies the correlation between recipient genotype and cooperative actor genotype^58–60^, must exceed the cost of the cooperative behavior^18^. Putting this principle into the context of our experiments, the cost of RbmC and Bap1 production is set by their regulation (here, nearly constitutive) and nutrient supply conditions, and the benefit is set by the extent to which shear stress is applied to biofilms (Fig. S8). The relatedness coefficient – taking note again that this effectively measures the extent to which RbmC/Bap1-producing cells benefit each other relative to the total population composition^58,60–63^ – is controlled by the range over which RbmC and Bap1 diffuse and the spatial distribution of producer and cheater cell clusters^26,28,29,58,63,64^. In many reported cases, the clustering of clonal cell lineages within mixed biofilm communities can be sufficient to stabilize cooperation against cheating^25–29^. Here, however, the spatial range of RbmC/Bap1 sharing is larger than the size of clonal cell clusters due to the leakage of these proteins from groups of producing cells, and the associated public goods dilemma cannot be resolved by clonal group clustering alone. Instead, the solution is rooted in the combination of biophysical mechanisms of protein retention and diffusion, community spatial structure, and environmental perturbations, all captured in the spatial exploitation model (Fig. 3).

The key parameter in our model, the exploitation radius *R*, can additionally depend on many physiological factors including the production rate of the matrix, duration of biofilm growth, aggregation of adhesion proteins, uptake of adhesion protein by cheater matrix, and biofilm dispersal^65^, among other factors. Competition for binding to the exopolysaccharides and for surface adsorption from other biomacromolecules in the environment may also reduce *R*^66^. In addition, although we have assumed *R* to be constant in our spatial model of *P*_protection_, *R* does appear to vary and drop to a much smaller value when the inoculation number density goes above 10^−1^ cells/100 μm^2^ (Fig. S9). This can be understood as follows: At a high total inoculation number density, the size of each producer cluster is smaller due to nutrient limitation and the concentration of secreted proteins around each producer cluster is consequently reduced. Additionally, the elevated incorporation of adhesion proteins into the surrounding cheater clusters further constrains the exploitation range. In the limit of a confluent biofilm layer, the system reduces to the well-studied case of a densely packed bacterial colony^21,23^. To account for the variation of *R* and *P*_protection_, a multiscale model that considers these additional factors should be implemented, which we leave for future studies. Nevertheless, our simplified spatial model already provides a good quantitative description of the underlying dynamics of cooperative matrix protein production, especially in the low seeding density regime most relevant to the natural conditions of *V. cholerae*. While the concept of exploitation range has been discussed in relation to public good production in bacterial communities^26–29,67–70^, direct quantitation of the exploitation radius and a close comparison between model and experiment, as we achieved here, represent important steps forward. Our study thus provides both a conceptual guideline and technical tool set for future studies on how to quantify public goods sharing in relation to population structure in the biofilm context.

Finally, previous work has shown that in the cases of extracellular enzymes^27,71,72^, siderophores^23,73^, and autoinducers^74–76^ as diffusive public goods, mostly studied in continuous films, clonal segregation and efficient consumption of public goods (or their enzymatic products and complexes) through high local cell density and a high uptake rate can minimize the potential for exploitation in the context of dense bacterial communities^21,26,27^. In the context of spatially discrete clonal clusters, we expect the evolutionary stability of these public goods to be determined by the interplay between the exploitation range and the spatial structure as shown here.

## Acknowledgments

We thank Andrew Sharo, Sara Siwiecki, and Carissa Chan for their help in the initial experiments. We thank Xin Huang, Japinder Nijjer and Qiuting Zhang for helpful discussions. Alexandre Persat, Kevin Foster, Benjamin Wucher, and James Winans provided helpful feedback on the manuscript. S.M. thanks Paul Fischer, Som Dutta, and Nek5000 user group for helpful discussions regarding numerical computations. Numerical computations were performed using the resources of the Minnesota Supercomputing Institute at the University of Minnesota. J.Y. holds a Career Award at the Scientific Interface from the Burroughs Wellcome Fund. C.D.N. is supported by the Simons Foundation award number 826672. J.T. holds a Career Award at the Scientific Interface from the Burroughs Wellcome Fund. J.-S.B.T. is a Damon Runyon Fellow supported by the Damon Runyon Cancer Research Foundation (DRG-2446-21). Research reported in this publication was supported by the National Institute of General Medical Sciences of the National Institutes of Health under Award Number DP2GM146253 (awarded to J.Y.).

## Author contributions

C.D.N. and J.Y. conceived the project. J.-S.B.T. and T.N. performed the experiments. J.Y. performed strain cloning. S.M and J.T. performed fluid dynamics modeling. J.-S.B.T. performed ecological modeling. J.-S.B.T., S.M., J.T., C.D.N. and J.Y. analyzed the data. All authors contributed to the writing of the paper.

## Competition interests

The authors declare no competing interests.

## Methods

### Strains and media

All *V. cholerae* strains used in this study are derivatives of the wild-type *Vibrio cholerae* O1 biovar El Tor strain C6706 and listed in Table S1. All strains harbor a missense mutation in the *vpvC* gene (*vpvC*^W240R^) that elevates intracellular c-di-GMP level and are robust biofilm formers^44^. This allows us to focus on the ecological consequences of adhesion protein production in *V. cholerae* biofilms rather than behaviors involving gene regulation. Additional mutations were genetically engineered using the natural transformation method or through the suicide vector pKAS32^77,78^. All strains were grown overnight in lysogeny broth (LB) at 37°C with shaking. Competition assays were performed in M9 medium and supplemented with 2 mM MgSO_4_, 100 μM CaCl_2_, 0.5% glucose, and 0.1 mg/mL of bovine serum albumin (BSA). BSA is required for blocking non-specific surface adsorption in the immunostaining experiments; to be consistent, we include BSA in all competition assays as well.

### Protein quantification

*V. cholerae* strains encoding Bap1-3×FLAG or RbmC-3×FLAG were grown in culture tubes containing 3 mL LB and sterile glass beads overnight at 30°C with shaking. The next day, cultures were vortexed to break up pellicles and cell clusters and the OD_600_ was measured. 1 mL of cell suspensions were transferred to sterile 1.5 mL microcentrifuge tube and spun at 18,000g for 3 min. 500 μL of the cell supernatant were transferred to a fresh 1.5 mL microcentrifuge tube and the rest was discarded from the pellet. The cell pellets were lysed for 30 min in 100 μL lysis solution [1× Bugbuster (EMD Millipore 70921), lysozyme (0.1 mg/mL), and benzonase™ nuclease (Sigma E1014)] and then brought to a final volume of 1 mL with 1× PBS. 30 μL of each cell suspension was combined with 10 μL of 4× SDS PAGE sample buffer (40% Glycerol, 240 mM Tris pH6.8, 8% SDS, 0.04% Bromophenol Blue, 5% β-mercaptoethanol) and boiled for 10 minutes at 95°C. Samples were run on a 4-15% Mini-PROTEAN TGX gel (BioRad 4568086) in 1× SDS PAGE running buffer (25 mM Tris, 192 mM Glycine, 1% SDS, pH 8.3) at 120 V for 70 minutes at 4°C. The proteins were transferred to a PVDF membrane (BioRad 1620174) in 1×transfer buffer (25 mM Tris, 192 mM Glycine, 10% methanol, pH 8.3) at 100V for 1 hour at 4°C. Membranes were incubated in 5% milk in TBST overnight at 4°C then at room temperature for 1hr. Following incubation, membranes were washed 3 × 10 minutes in 1× TBST (American Bio AB14330-01000). The membranes were blotted using α-DYKDDDDK at 0.1 μg/mL (Biolegend 637311) in 1× TBST with 3% BSA for 1hr at room temperature and washed 3 × 10 minutes with 1× TBST. Blots were developed using the Super Signal PLUS Pico West Chemiluminescent Substrate (Thermofisher 34580) for 5 min and pictures taken using a BioRad Chemidoc-MP. Analyses of sample signal were performed in ImageJ. The total signal for whole cell lysate (Pellet) and supernatant (Sup.) were measured by densitometry and each fraction was calculated against the total signal.

### Static competition assay with end-point flow disturbance

*V. cholerae* strains carrying different fluorescent proteins were streaked onto an LB agar plate and grown overnight at 37°C. Single colonies with a rugose morphology were picked and grown overnight at 37°C with glass beads. The LB culture was then vortexed vigorously before back-diluting 30-fold in M9 medium and grown for 3-4 h with shaking and with glass beads. After another vortexing, the inoculant was washed with fresh M9 medium without glucose 3 times. The inoculant was then bead bashed using a Digital Disrupter Genie with small glass beads (Acid-washed, 425–500 μm, Sigma). This procedure ensures that large cell clusters formed in culture were broken apart to allow more accurate measurement of OD_600_. The inoculant of the two competing strains was then mixed at different ratios and OD_600_, and 100 μL of the mixture was added to wells of 96-plates with a #1.5 coverslip bottom (MatTek). The cells were allowed 1h to attach, after which the wells were washed twice with fresh M9 medium. The wells were subsequently filled with 100 μL M9 growth medium with glucose and 0.1 mg/mL of BSA and imaged by a confocal fluorescence microscope (see below) to quantify the inoculation number density for both strains. Inoculation number densities, instead of OD_600_, were used to quantify *f*_0,p_ due to the asymmetry in inoculation efficiency between producer and cheater (Fig. S10). The cultures were then grown for 16-18 h at 30°C before biomass quantification. After growth, one set of biomass quantification for both strains was done by fluorescence confocal microscopy before a flow disturbance to the culture was introduced by washing the well vigorously twice with fresh medium, after which another set of biomass quantification was done. Immunostaining of adhesion proteins was done using producer strains expressing C-terminal 3×FLAG-tagged RbmC or Bap1 co-cultured with the cheater under the same growth conditions for competition assays, except that the growth medium contained Cy3-conjugated anti-FLAG antibodies (2 μg/mL; Sigma-Aldrich).

### Competition assay under flow

The inoculants of the competition strains were prepared in the same way as described in the static competition assay. The inoculants were then introduced into microfluidic channels (channel dimensions: 1 cm in length, 400 μm in width and 60 μm in height) through the outlet without inoculating the inlet. This ensures that no biofilms were grown in the zero-flow zone directly beneath the inlet. The cells were allowed 2h to attach, after which sterile inlet and outlet polytetrafluoroethylene tubing were connected to the microfluidic chamber and the M9 growth medium was flown through the channel at flow rates controlled by a syringe pump. The flow rates used in this study range from 0.1 to 1 μL/min, corresponding to average flow speeds from 69.4 to 694 μm/s and shear rates from 7 to 70 s^−1^. The microfluidic channels were transferred to a confocal fluorescence microscope and imaged to quantify the inoculation number density for both strains. The cultures under flow were then grown for 16-18 h at 30°C before biomass quantification. After growth, one set of biomass quantification for both strains were done by fluorescence microscopy.

### Microscopy, biomass quantification, and growth rate measurement

Fluorescence confocal microscopy was conducted using a Yokogawa W1 confocal scanner unit connected to a Nikon Ti2-E inverted microscope with a Perfect Focus System. Cells constitutively expressing SCFP3A, mNeonGreen, and mScarlet-I cytosolically were excited by lasers at 445 nm, 488 nm, and 561 nm, respectively, and the fluorescent signal was recorded by a sCMOS camera (Photometrics Prime BSI) after the corresponding filters. Confocal images were taken using a 60× water immersion objective (NA = 1.2) at 2×2 pixel binning and the resulting pixel size was 217 nm. All images presented are raw images rendered with Nikon NIS-Elements. Producers are pseudo-colored red, cheaters are pseudo-colored yellow, and the immunosignals are pseudo-colored cyan throughout the text unless indicated otherwise.

To quantify the inoculation number density before growth, an image at the surface was taken for each wavelength at the center of each well or microfluidic channel. To quantify the biomass after growth, a 3D tiled image with a 3 μm step size in *z* was taken at 3-5 evenly distributed positions within same the area for each inoculation number density. The total height of the 3D image was chosen to include all biomass of the surface-attached biofilm and is 60-72 μm in the static competition assay and 60 μm for the competition assay under flow.

For accurate segmentation and quantification of the biomass, we used a modified Bernsen’s local thresholding method which is less sensitive to depth-dependent intensity variations and contrast reduction (due to pinhole crosstalk and scattering) in the 3D confocal image. Briefly, each 2D image slice first went through a constant background subtraction and denoising, and the image contrast was enhanced by subtracting the image by its Gaussian blur. To determine whether a pixel belongs to cells or background, if the contrast in the local neighborhood around a given pixel is higher than the contrast threshold (50 in all data set), the pixel is set to belong to cells if its value is higher than 1.5 times the median value of the neighborhood; otherwise a pixel is set to belong to cells if its value is higher than an intensity threshold determined based on the brightness of the image. Examples of segmented images are shown in Fig. S4. The total area of cells in each 2D image was converted to a number density using the average cell sizes, measured separately at low cell densities for different fluorescence channels. The number densities in each 2D slice were then added to obtain the total 3D biomass.

Growth rates of producer and cheater biofilms were measured by time-course imaging of biofilm growing in 96-well plate. Confocal images were taken every 2 h for 40 h and biomass was quantified as described above at each time point. The effect of confocal imaging was confirmed to be nonsignificant to biofilm growth under the conditions used in the experiment. We extracted growth rates as slopes of fitting lines to the growth curves in semi-log plots in their exponential phase. Carrying capacities used in the competition model were measured as the maximum biomass in the stationary phase.

### Quantification of exploitation range and protection probability

Cross-correlations between producer and cheater biomass distributions were used to quantify the exploitation radius *R*. We model the cheater biomass distribution as being exponentially decaying around a producer cell cluster of radius *r*_0_ with the decaying constant equal to *R* - *r*_0_; this yields an exploitation radius *R*. We found that the cross-correlation between the producer and cheater images gives adequate information to extract information on *r*_0_ and *R* (Fig. S6). To obtain cross-correlations from experimental images, inoculant mixtures of producer strain with OD_600_ = 0.001 and cheater strain with OD_600_ = 0.1 were used for static competition assay with isolated producer clusters. Maximum intensity projections of confocal images for both strains after end-point flow disturbance were thresholded and used for cross-correlation calculations. The field of view of each image is ~1.8 mm × 1.8 mm. The cross-correlation between producer and cheater was calculated as

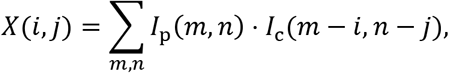

where *I*_p_ and *I*_c_ are the binary images for producer and cheater, and *m* and *n* are the row and column indices of the image. *X*(*i*, *j*) was then radially averaged and converted to real spatial units. The error in Fig. 3C was calculated as the standard error from the N = 4 replicates. For a negative control of the correlated biomass distribution, we compared the normalized cross-correlation between producer and cheater biofilms with that between producer and producer biofilms (Fig. S6). The normalized cross-correlation is defined as

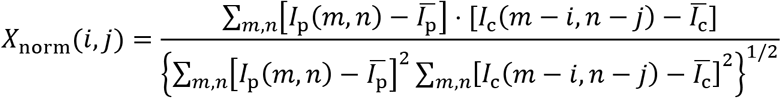

Where 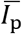 and 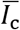 are the mean values of *I*_p_ and *I*_c_, respectively.

### Spatial exploitation model based on randomly distributed disks

Assuming each producer cluster contributes a circular disk of protection with area *πR*^2^, the overall protection probability conferred by producer biofilms to cheater biofilms in an area of interest can be modeled by the coverage problem of randomly distributed circular disks^79^: let there be *n* interpenetrable disks with area *a* distributed randomly in an area of interest *A*, the uncovered probability of *A* has a mean value 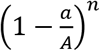. In the limit of large *A* and 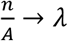, the uncovered probability approaches exp (−*λa*). Therefore, for producer clusters with number density *σ*_0,p_ and exploitation radius *R*, the protection probability reads *P*_protection_ = 1 − exp(−*σ*_0,p_ ⋅ *πR*^2^).

### Flow and adhesion protein simulation in a microfluidic environment

To gain insight into the effect of different fluid flow speeds on the transport of molecules released by the producer, we solve the governing equations of the fluid flow and a coupled scalar transport equation to model the molecular distribution around a producer. We assume the producer cluster is shaped like a solid hemisphere adhered to the bottom surface of a rectangular channel and subjected to fluid flow, as shown in Fig. S11. The fluid flow is governed by the Navier-Stokes and continuity equations, and the molecular transport is described by the coupled advection-diffusion equation, which are respectively given by:

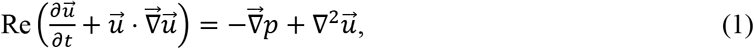

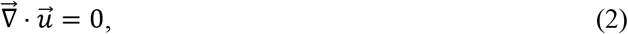

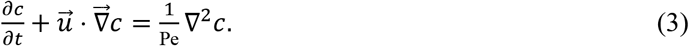

In these equations, 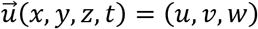 is the fluid velocity vector, *p*(*x*, *y*, *z*, *t*) is the pressure, and *c*(*x*, *y*, *z*, *t*) is the concentration of molecules released by the producer. Eqs. (1-3) are nondimensionalized using a length scale *d* = 20 μm, which is the average diameter of a producer cluster, as found in experiments. The time scale used is the advective time scale 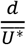 where 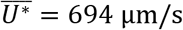 is the dimensional average inflow velocity. The concentration is normalized using an arbitrary reference concentration of *c*_0_ = 1 adhesion molecule/μm^3^.

There are two important nondimensional parameters that describe the equations. The Reynolds number 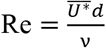 is defined as the ratio of the inertial to the viscous forces, where ν = 1 x 10^−6^m^2^/s is the kinematic viscosity of water. Our experiments are typically in the range 0.01 < Re < 0.1, meaning viscous effects dominate the dynamics. At these Re, the left-hand side of Eq. (1) can be neglected, reducing to the Stokes equation. The Péclet number, 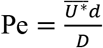 is defined as the ratio of the diffusive time scale to the advective time scale; *D* = 80 μm^2^/s is the diffusion coefficient of the adhesion molecules based on the Stokes-Einstein equation and their estimated size. Pe > 1 suggests fluid advection has a larger role in transporting the molecules than diffusive transport. We vary the Péclet number in the range 0 ≤ Pe ≤ 173.5, in agreement with variable experimental conditions of 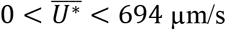.

We simulate fluid flow through a channel of dimension *L_x_* × *L_y_* × *L_z_* as shown in Fig. S11. The solid hemisphere modeling the producer cluster is positioned at 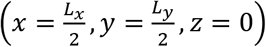. In the experiments, the aspect ratio of the channel is Γ_*x*_ × Γ_*y*_ × Γ_*z*_ = 500 × 20 × 3, where 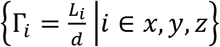. This large aspect ratio makes simulations computationally expensive, so we instead use a channel of dimensions Γ_*x*_ × Γ_*y*_ × Γ_*z*_ = 20 × 6 × 3 in the simulation, which preserves the important cross-stream dimension along the z-axis.

A no-slip boundary condition is imposed for the top and bottom surfaces of the domain 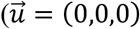 at *z* = {0, *L_z_*}). A Dirichlet boundary condition is applied at the inlet 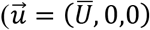 at *x* = 0), where 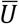 is the average nondimensional inflow velocity into the channel scaled by 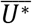 and 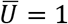 corresponds to an average velocity of 694 μm/s. An open/outflow boundary condition is applied to the outlet at *x* = *L_x_*. Along the material walls at *y* = {0, *L_y_*}, we impose a free-slip or no-penetration boundary condition. The boundary condition for the concentration is no-flux on all material walls, except for the surface of the solid hemisphere, for which we use a time-varying Dirichlet boundary condition of the form,

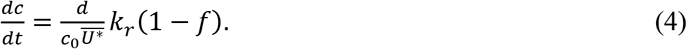

Here, *k_r_* = 0.015 molecules/μm^3^/s is the total production rate of the molecules from the surface of the producer cluster^43^ and *f* quantifies the fraction of molecules retained in the producer cluster based on experimentally measured values. Equation (4) suggests that there is a constant rate of production of molecules at the surface of the hemisphere.

We numerically solve the governing equations (1-4) using the spectral element fluid solver NEK5000^80^. NEK5000 has been used to study a broad range of fluid flow problems^81^, including complicated chaotic fluid flows and reaction-advection-diffusion systems^82–84^. Our approach uses a semi-implicit operator-splitting scheme which is 2^nd^ order accurate in time and exponentially convergent in space. Our domain consists of 392 hexahedral spectral elements, and we have used a 17^th^ order Lagrangian interpolant polynomial for spatial discretization, which we found is sufficient to resolve both the fluid flow and the intricate features of the molecular distribution around the producer.

To resolve the small Re of the flow, we evolve Eqs. (1-2) using an extremely small time step of Δ*t* = 1 x 10^−8^. We simulate the flow field for 1.4 x 10^6^ steps and ensure that the flow field reaches a steady state. We then stop the evolution of the fluid flow and use the steady velocity profile to solve the advection-diffusion equation (3). Since we do not evolve the flow field, we use larger time steps (typically Δ*t* = 1 x 10^−3^) to evolve the concentration field for a total of 3 min (Fig. S12).

### Statistics

Errors correspond to SEs from measurements taken from distinct samples unless mentioned otherwise. Standard *t*-tests were used to compare treatment groups and are indicated in each figure caption. Tests were always two-tailed and unpaired as demanded by the details of the experimental design. All statistical analyses were performed using GraphPad Prism software.

