## Supplementary information for "Social evolution of shared biofilm matrix components"

#### Supplementary Methods

##### Two-part competition model in a static environment

Competition between adhesion protein producer and cheater in a static environment before and after washing was modeled using a two-part model: 1) Structureless competition model of co-cultured producer and cheater in a static environment and 2) Spatial model of exploitation capturing the effect of disturbance introduced by washing. In the first part, we use the classic Lotka-Volterra model to model co-cultured producer and cheater populations competing for the same nutrient source[1]. The populations for producer and cheater cells, $P_{\mathrm{p}}$ and $P_{\mathrm{c}}$, both follow the logistic growth function, but with different growth rates $r_{\mathrm{p}}$ and $r_{\mathrm{c}}$ measured separately from their growth curves in their mono-cultures (Fig. 1a). The carrying capacity for producer and cheater of the environment are $N_{\mathrm{p}}$ and $N_{\mathrm{c}}$. The equations read

$$\begin{cases} \dfrac{dP_{\mathrm{p}}}{dt} = r_{\mathrm{p}}P_{\mathrm{p}}(1 - \dfrac{P_{\mathrm{p}} + \alpha_{\mathrm{pc}}P_{\mathrm{c}}}{N_{\mathrm{p}}}) \\ \dfrac{dP_{\mathrm{c}}}{dt} = r_{\mathrm{c}}P_{\mathrm{c}}(1 - \dfrac{P_{\mathrm{c}} + \alpha_{\mathrm{cp}}P_{\mathrm{p}}}{N_{\mathrm{c}}}) \end{cases}.$$

Here $\alpha_{\mathrm{pc}}$ and $\alpha_{\mathrm{cp}}$ represent the inter-strain effects and are both set to 1, since the competing producer and cheater strains have similar effects on each other in terms of nutrient depletion. The carrying capacity $N_{\mathrm{p}}$ and $N_{\mathrm{c}}$ specify the maximum populations the environment can sustain for each strain. In the Lotka-Volterra model, when $N_{\mathrm{p}} \neq N_{\mathrm{c}}$, the effective growth rate can become negative for the strain with lower capacity and the population of the strain with higher carrying capacity grows at the expense of the population of the low-capacity strain. Since we did not observe a reduction in population in either of the strains in the experiment, we set $N_{\mathrm{p}} = N_{\mathrm{c}}$ to be the maximum population of producer measured in the stationary phase (~40 h) to ensure that the effective growth rates always stay positive. The equations were numerically solved using the ode45 solver in MATLAB at 16 h for different initial producer frequencies $f_{0,\mathrm{p}}$ and inoculation number densities $\sigma_0$, and the frequency change before washing is defined as $\Delta f_{1,\mathrm{p}} \equiv P_{\mathrm{p}}/(P_{\mathrm{p}} + P_{\mathrm{c}}) - f_{0,\mathrm{p}}$. The results are shown in Fig. S5a.

The second part of the model uses the spatial exploitation model discussed in the main text and Methods section. After washing, all $P_{\mathrm{p}}$ remained adherent to the surface, while only $P_{\mathrm{protection}} = 1 - \exp(-\sigma_{0,\mathrm{p}} \cdot \pi R^2)$ of $P_{\mathrm{c}}$ remained adherent as a result of scavenging adhesion proteins from the producer. Therefore, the frequency change after washing was calculated as $\Delta f_{2,\mathrm{p}} \equiv P_{\mathrm{p}}/(P_{\mathrm{p}} + P_{\mathrm{protection}} \cdot P_{\mathrm{c}}) - f_{0,\mathrm{p}}$. Here $\sigma_{0,\mathrm{p}} = f_{0,\mathrm{p}}\sigma_0$. The results are shown in Fig. S5b. The numerical results agree well with the experimental data measured both before and after washing (Fig. 2d). The parameters used in the two-part competition model are summarized in Table S2. All parameters used in the modeling were experimentally calibrated.

#### Supplementary Figures

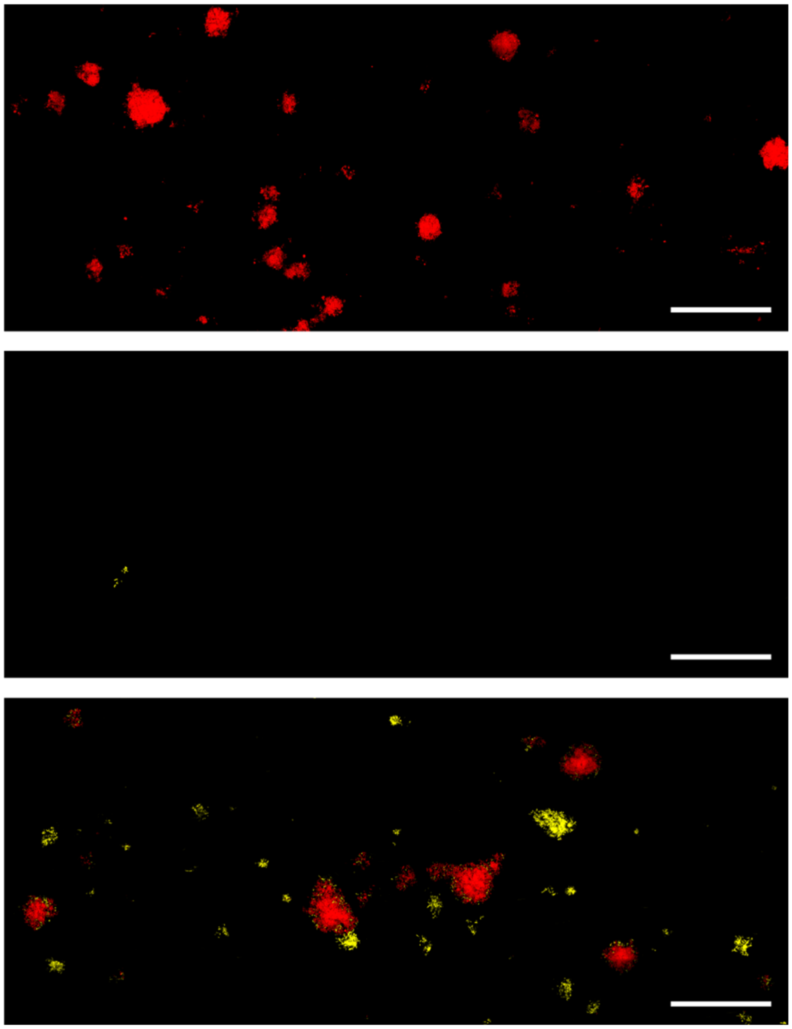

Fig. S1 | Adhesion protein sharing and exploitation between producer and cheater in wild-type (WT) *V. cholerae*. Confocal images at 6 µm away from substrata of WT adhesion protein producer (red) and cheater (yellow) in their mono-cultures (*top* and *middle*) and co-cultures (*bottom*) in a flow environment. These data show that exploitation of adhesion proteins by cheaters is not limited to strains in a constitutive biofilm producing background, but generalizable to WT strains in which biofilm formation is regulated. Scale bars: 100 µm. Experiments were performed under flow as described in the main text at a flow rate of 0.6 µL/min and in M9 growth medium supplemented with 0.5% glucose and 0.5% casamino acids (Difco Laboratories).

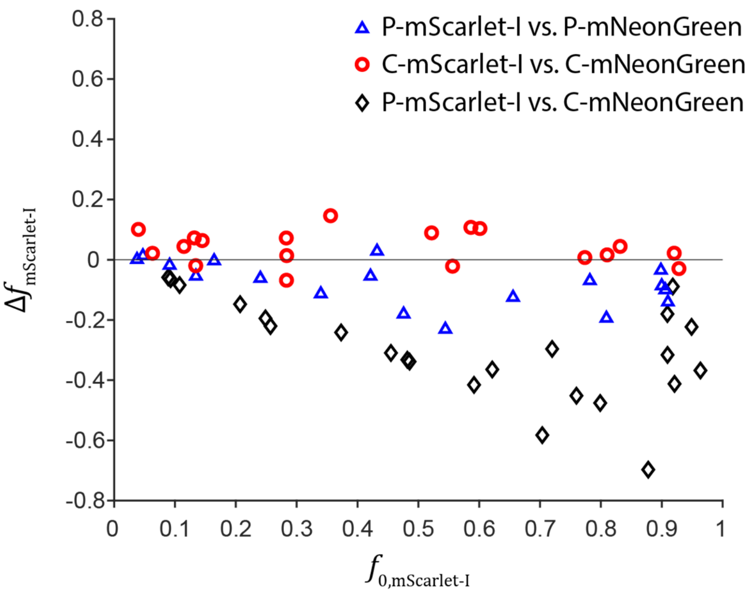


Fig. S2 | Competition between strains expressing different fluorescent proteins in a static environment with no washing. The frequency change is more significant for competition between producer and cheater strains (abbreviated as P and C, respectively; P-mScarlet-I vs. C-mNeonGreen) than between isogenic strains expressing different fluorescent proteins (P-mScarlet-I vs. P-mNeonGreen and C-mScarlet-I vs. C-mNeonGreen). We do note a slight growth advantage of P-mNeonGreen over P-mScarlet-I, which could be due to the slightly higher metabolic cost associated with the production of the red fluorescent protein.

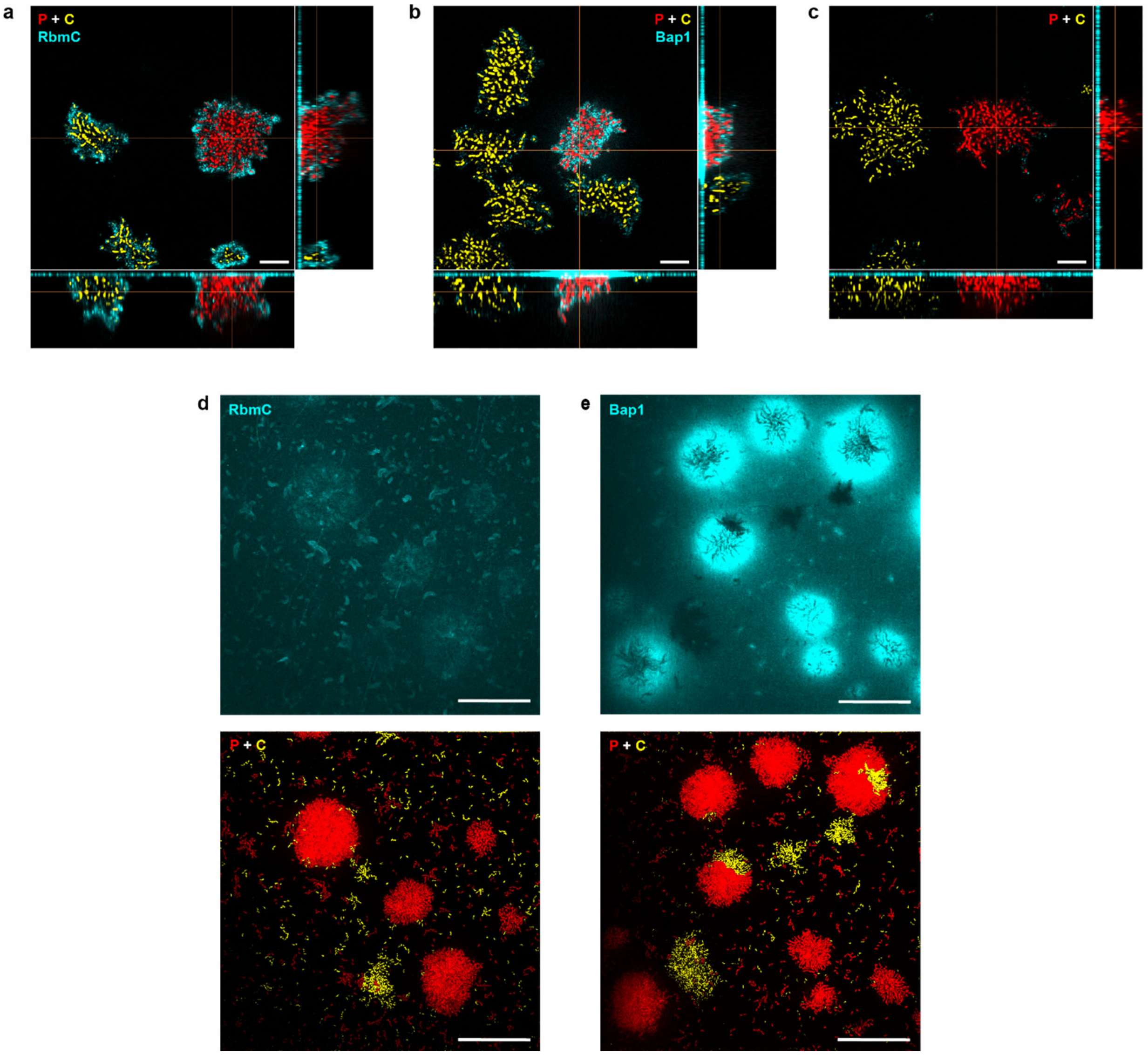


Fig. S3 | Distribution of adhesion proteins among producer (abbreviated as P, red) and cheater (abbreviated as C, yellow) biofilms shown by Cy3-conjugated anti-FLAG antibody staining (cyan). **a-c**, Orthogonal views of a producer strain carrying 3×FLAG-tagged RbmC (a) and a producer strain carrying 3×FLAG-tagged Bap1 (b) co-cultured with the cheater. The epitopes are on the C-termini. A control of co-cultured producer (non-FLAG-tagged) and cheater under the same staining condition is shown in (c). Scale bars: 10 μm. **d-e**, Surface signal in a different set of images of a producer strain carrying 3×FLAG-tagged RbmC (d) and a producer strain carrying 3×FLAG-tagged Bap1 (e) co-cultured with the cheater. Scale bars: 50 μm.

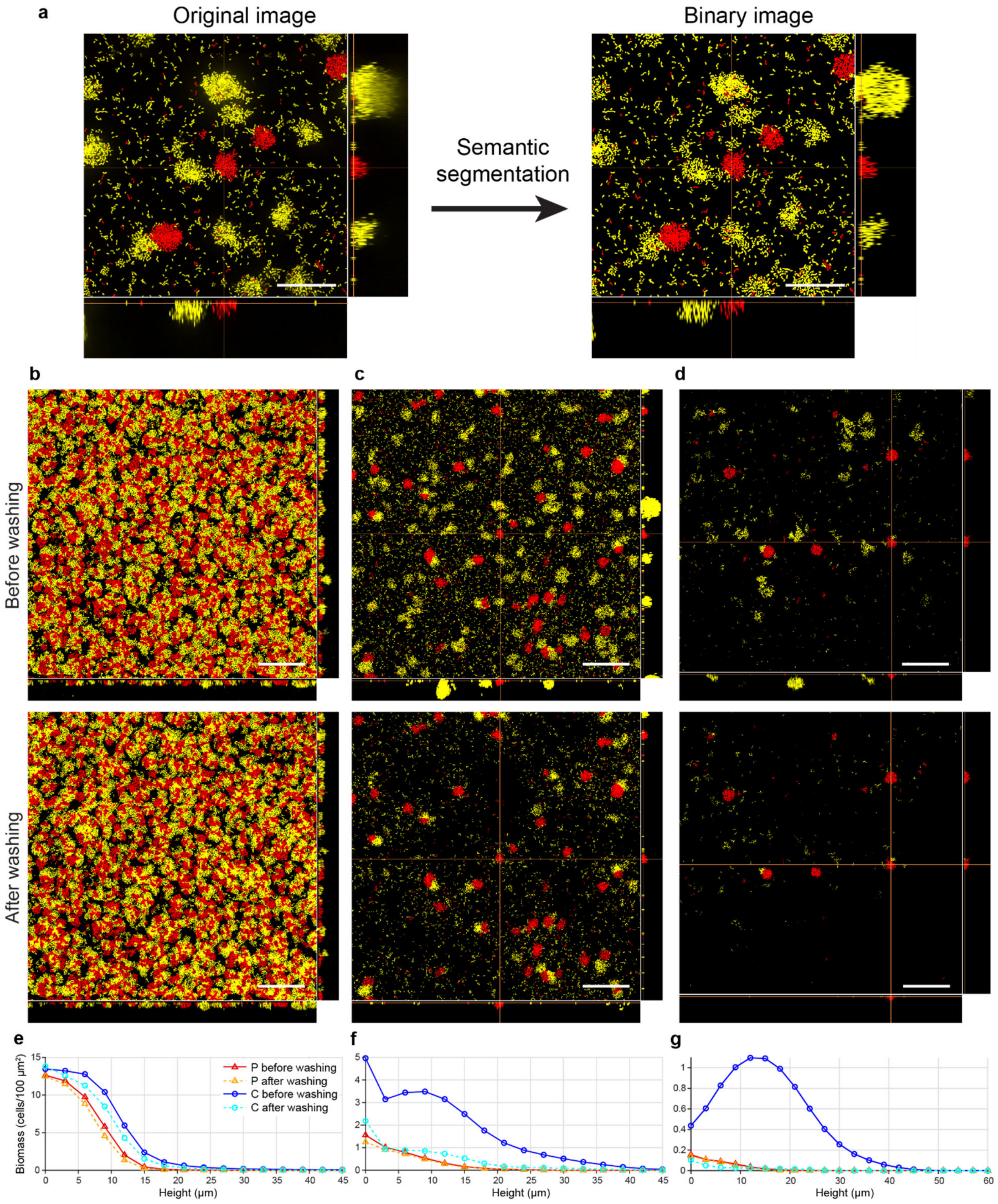


Fig. S4 | Image analysis and biomass quantification. **a**, Original 3D fluorescence image (*Left*) was segmented by a local thresholding method to obtain a 3D binary image for both producer and cheater cells for biomass quantification (*Right*). Scale bars: 50 µm. **b-d**, Segmented binary images corresponding to Fig. 2a-c. Scale bars: 100 µm. **e-g**, Layer-by-layer biomass quantification of binary images in (b-d). The biomass data show that the majority of biomass in producer biofilms remain adherent after washing, regardless of the density of cells, while the adherence of cheater biofilms after washing depends strongly on the density of the producer. The biomass distributions also show distinct differences along the dimension vertical to the substratum: at medium and low density (f and g), the producer biomass is localized near the substratum and peaks at the surface, while the cheater biomass is distributed further away from the substratum. This may provide additional fitness advantages to the producer in the presence of a nutrient gradient near the solid substratum, a common scenario for microbes in marine and fresh-water habitats. After washing, the biomass distributions for producers and cheaters are both localized at the surface, suggesting correlations among adhesion protein sharing, biofilm morphology, and adhesion.

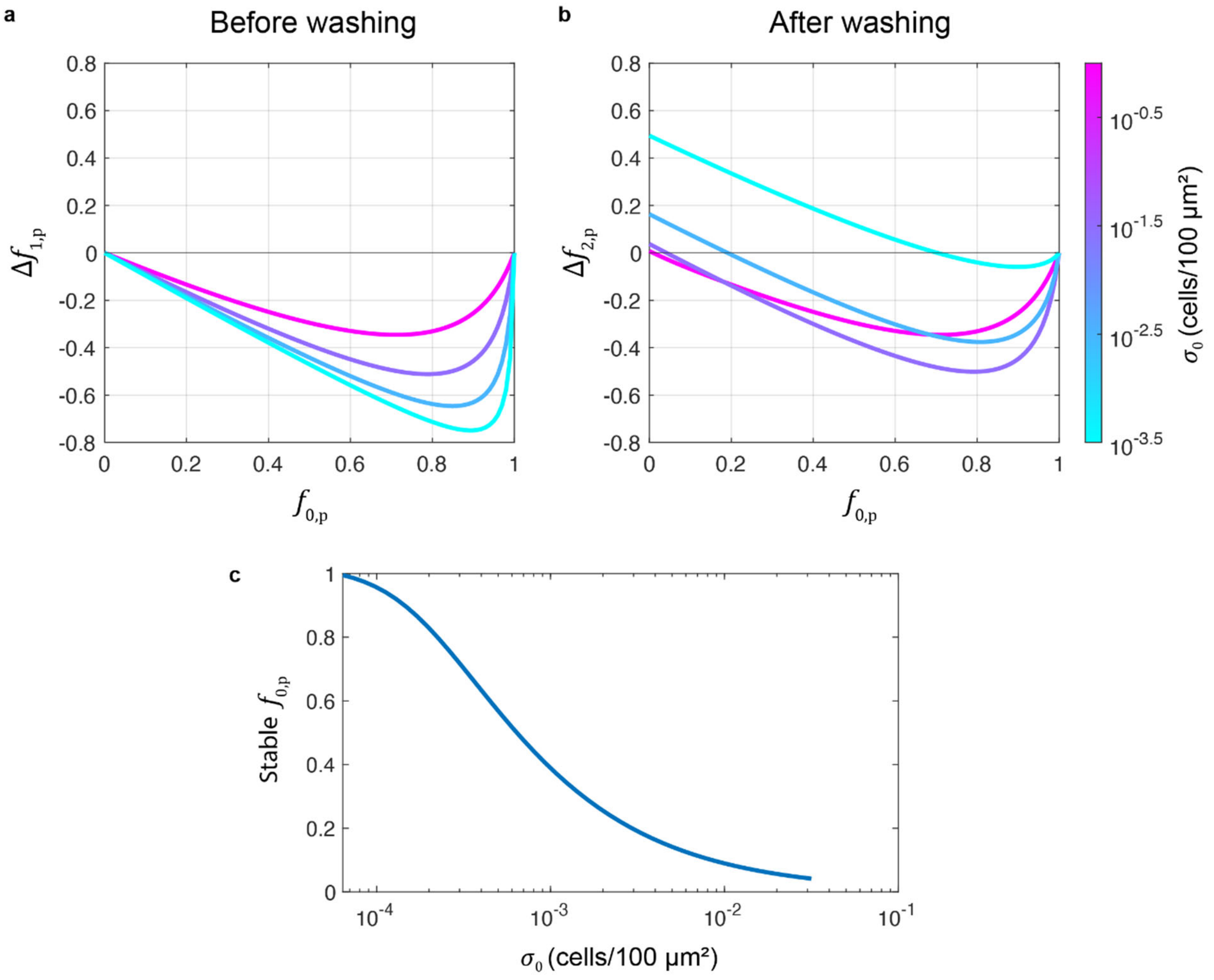


Fig. S5 | A two-part competition model between adhesion protein producer and cheater reproduces the experimental competition data. **a-b**, Competition between adhesion protein producer and cheater in a static environment before and after washing was modeled using a two-part model: 1) Structureless competition model of co-cultured producer and cheater in a static environment (*Left*) and 2) Spatial model of exploitation capturing the effect of disturbance introduced by washing (*Right*). See Supplementary Methods for more details. The numerical results agree well with the experimental data measured both before and after washing (Fig. 2d). In particular, before washing, the cheater outcompetes the producer and $\Delta f_{1,\mathrm{p}}$ decreases as $\sigma_0$ decreases. After washing, $\Delta f_{2,\mathrm{p}}$ is larger compared to $\Delta f_{1,\mathrm{p}}$, and the difference between them increases with decreasing $\sigma_0$. At intermediate $\sigma_0$, the negative frequency selection observed in experiments was reproduced. **c**, Stable $f_{0,\mathrm{p}}$ vs. $\sigma_0$ based on the two-part competition model. Our model predicts that the stable point moves from $f_{0,\mathrm{p}} = 0$ to $f_{0,\mathrm{p}} = 1$ as $\sigma_0$ decreases. Correspondingly, we predict that at very low inoculation densities, the producer always wins regardless of the initial frequency.

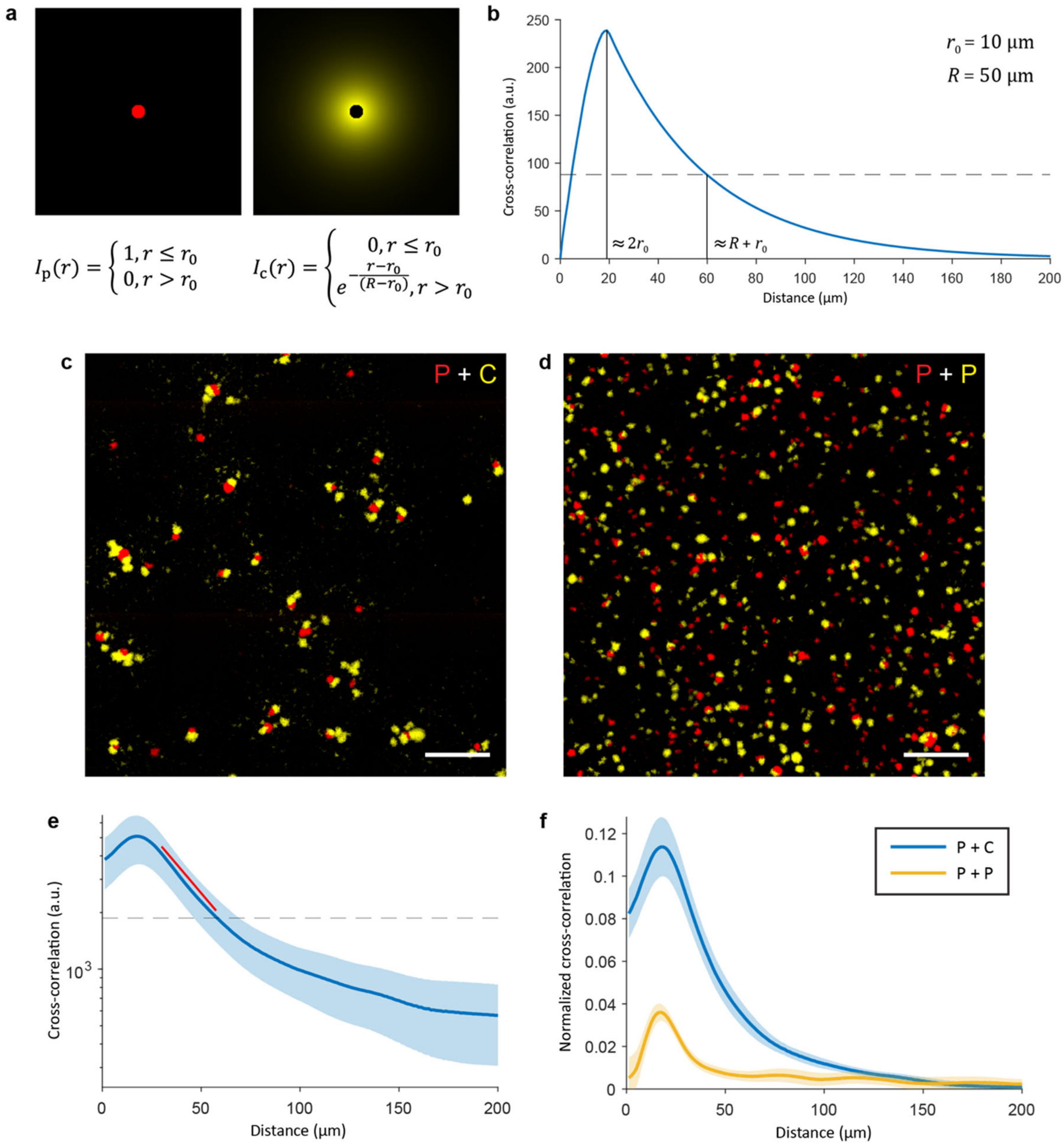


Fig. S6 | Quantification of exploitation radius through cross-correlation. **a**, Artificial images of a circular binary profile with radius $r_0$ mimicking the producer biomass distribution after wash (*Left*) and an exponentially decaying profile beyond $r_0$ with a width $R - r_0$ mimicking the cheater biomass distribution after washing (*Right*). The combined radius of exploitation is $R$. **b**, Cross-correlation between the artificial producer and cheater biofilm images in (a) shows a peak at $\approx 2r_0$ and a crossing with $e^{-1}$ of the peak value (dashed line) at $\approx R + r_0$. Here $r_0 = 10$ μm and $R = 50$ μm. **c-d**, Biomass distribution of co-cultured producer and cheater after washing (c) and co-cultured producers expressing different fluorescent proteins (d) as the control. Scale bars: 200 μm. **e**, Cross-correlation between producer and cheater images shows an exponentially decaying profile. The red line is a guide to the eye. **f**, Normalized cross-correlation between co-cultured producer and cheater after washing and between co-cultured producers expressing different fluorescent proteins as the negative control. The magnitude of the normalized cross-correlation in the former case is much larger, signifying the effect of protection. These data also show that the common peak at ~20 μm corresponds to the average diameter of the producer cluster.

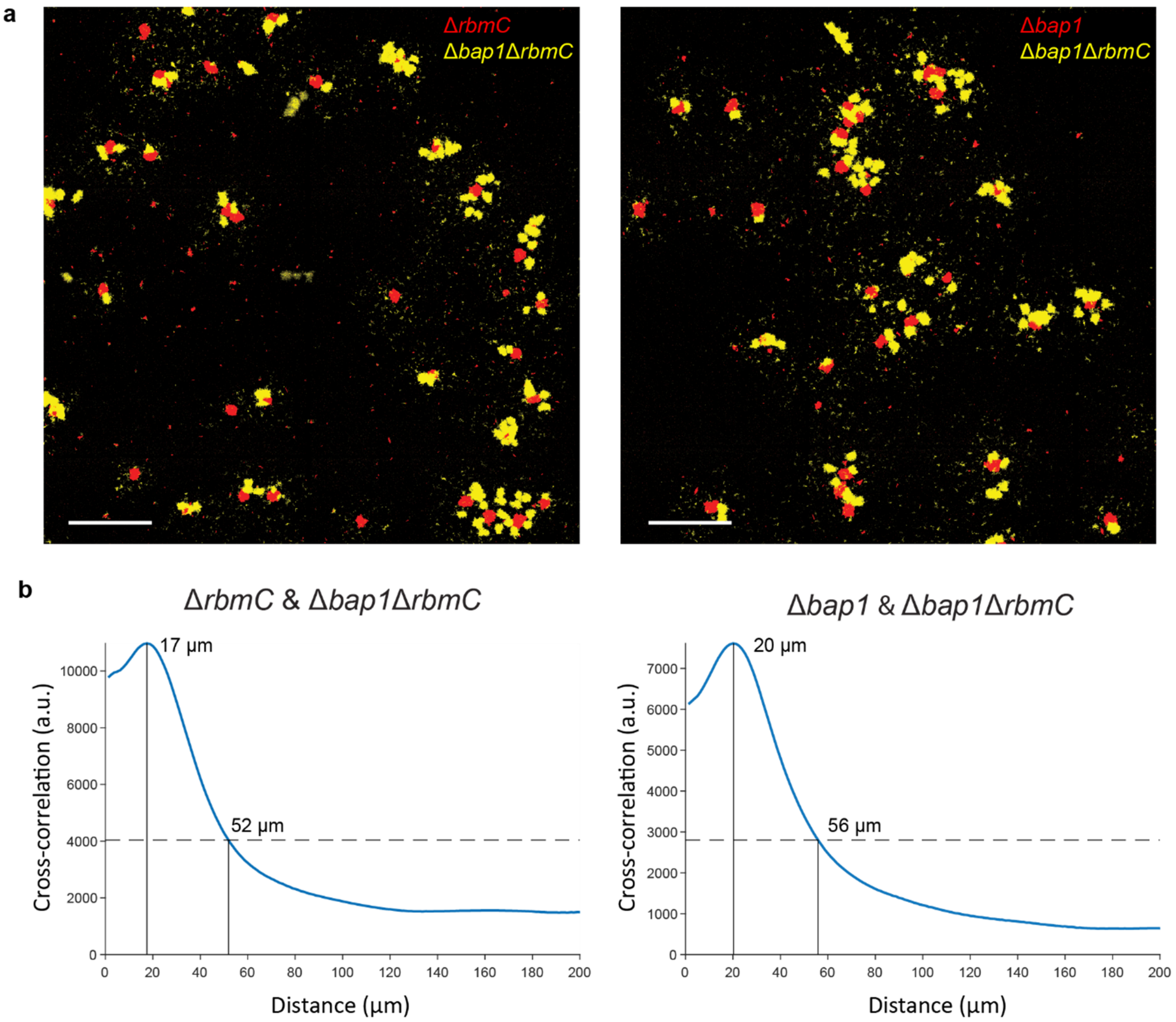


Fig. S7 | Bap1 and RbmC are redundant in the adhesion sharing assay. **a**, Images of co-cultured single adhesion protein mutants and double mutant after wash (*Left*: Δ*rbmC* & Δ*bap1*Δ*rbmC*, *Right*: Δ*bap1* & Δ*bap1*Δ*rbmC*). Scale bars: 200 μm. **b**, Cross-correlation between images of single mutant and double mutant biofilms. $R$ values extracted from the peak values and the crossing with $e^{-1}$ of the peak values (dashed lines) are 44 μm and 46 μm for the two cases, respectively. The single mutant biofilms adhere well to the surfaces, confirming the redundancy of the two adhesion proteins in attachment to glass surface. The similar exploitation radius, both slightly smaller than that conferred by the biofilms from the parental strain, shows that the two adhesion proteins are redundant and additive in the current competition assay, despite the different spatial distribution of RbmC and Bap1 in a biofilm (Fig. S3).

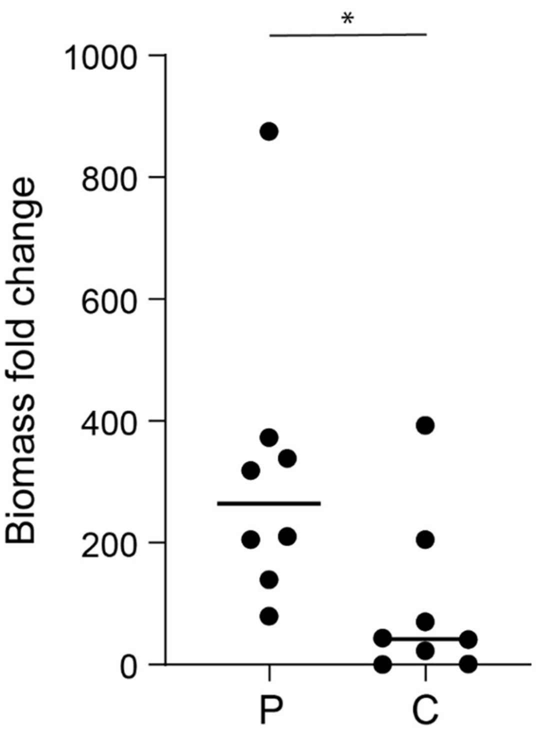


Fig. S8 | Biomass fold change of producer and cheater mono-cultures in a microfluidic flow chamber. Biomass fold change was measured after 16 h of growth in a flow chamber at a flow rate of 1 μL/min (*$P<0.05$, $N = 8$, Mann-Whitney test). These data show that, under a mild fluid shear, some cheater biofilms can remain adherent even without adhesion proteins. This is in contrast to the strong fluid shear induced by the washing step at the end of the static biofilm growth experiment in 96-well plate, where cheater biofilms were completely washed away in the absence of Bap1 and RbmC (Fig. 1 in main text). Therefore, the benefit of adhesion proteins to biofilm cells depends on the shear stress applied to them.

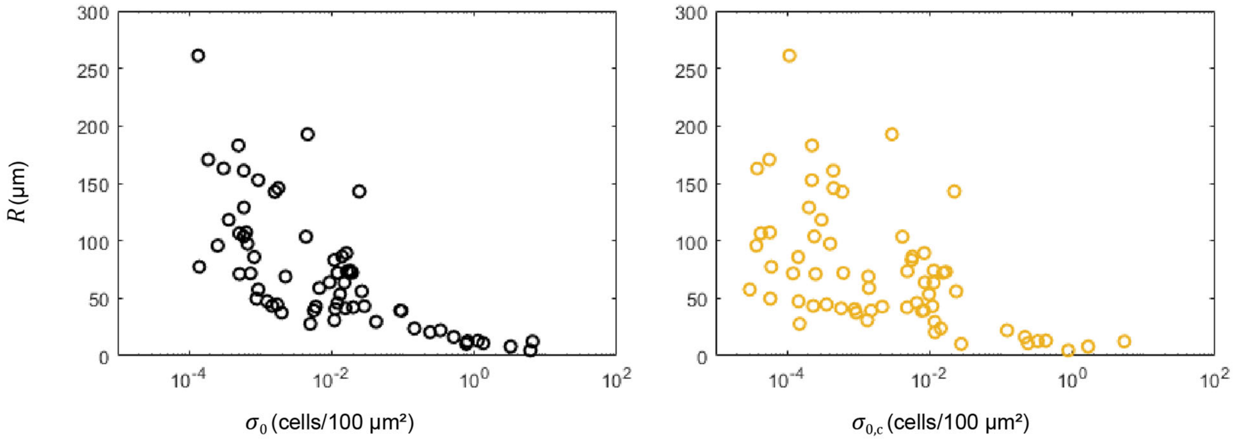


Fig. S9 | Dependence of $R$ on $\sigma_0$ and $\sigma_{0,c}$. Two regimes of $R$ are observed when $\sigma_0$ and $\sigma_{0,c}$ are below or above 0.1 cells/100 $\mu m^2$. At high $\sigma_0$, the size of each producer cluster and the amount of adhesion proteins secreted by each producer cluster are reduced, leading to a small $R$ (*Left*). On the other hand, at high $\sigma_{0,c}$, the binding of RbmC and Bap1 to the matrices in the surrounding cheater biofilms reduces the local adhesion protein concentration and therefore the exploitation radius $R$ (*Right*). Competition data from static competition assay in 96-well plate were used.

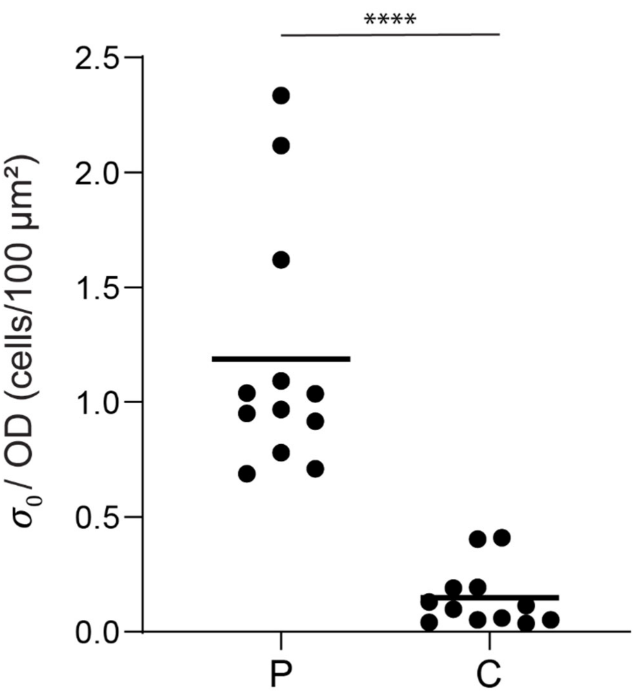


Fig. S10 | Inoculation efficiency of producer and cheater mono-cultures in a microfluidic flow chamber. Inoculation efficiency was defined as the ratio between initial surface coverage $\sigma_0$ and $OD_{600}$ of the inoculant (****P < 0.0001, N = 12, Mann-Whitney test). These data suggest that the cheater cells have a defect in initial colonization; therefore, for all of our biofilm quantifications we measured the inoculation surface densities ($\sigma_{0,\mathrm{p}}$ and $\sigma_{o,\mathrm{c}}$) to quantify $f_{0,\mathrm{p}}$ instead of using the $OD_{600}$ ratio in the liquid inoculants.

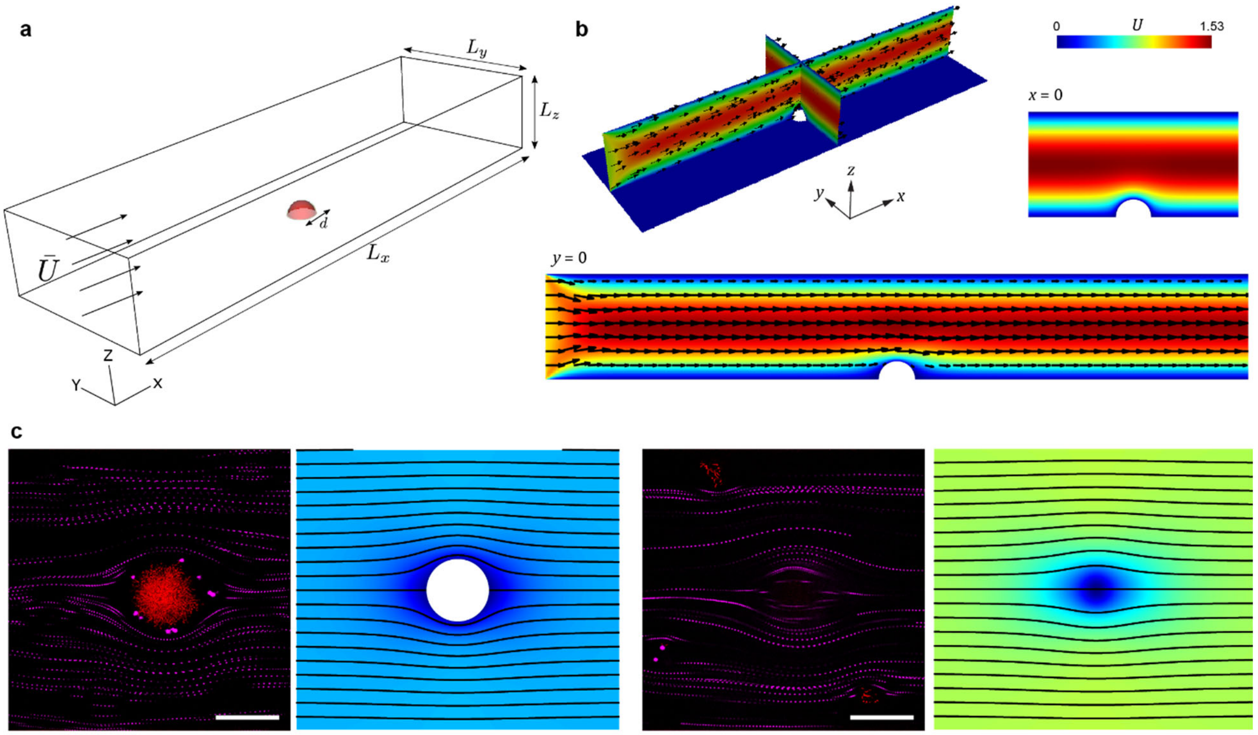


Fig. S11 | Simulated and experimental flow field around a producer cluster in a microfluidic chamber. **a**, Schematic of the computational domain. The producer cluster (red) was modeled as a hemisphere at the origin. **b**, The flow velocity field visualized by streamlines and colors around a producer cluster. **c**, Streamlines of flow imaged by 1μm fluorescent beads (magenta) suspended in the fluid around a producer cluster with diameter $r_0 \approx 40$ μm (red), 20 μm (left panel) and 42 μm (right panel; at the top of the cluster) away from the surface. The corresponding simulation results at $z = r_0/2$ and $z = r_0$ are shown on the right of each experimental image. Scale bars: 50 μm.

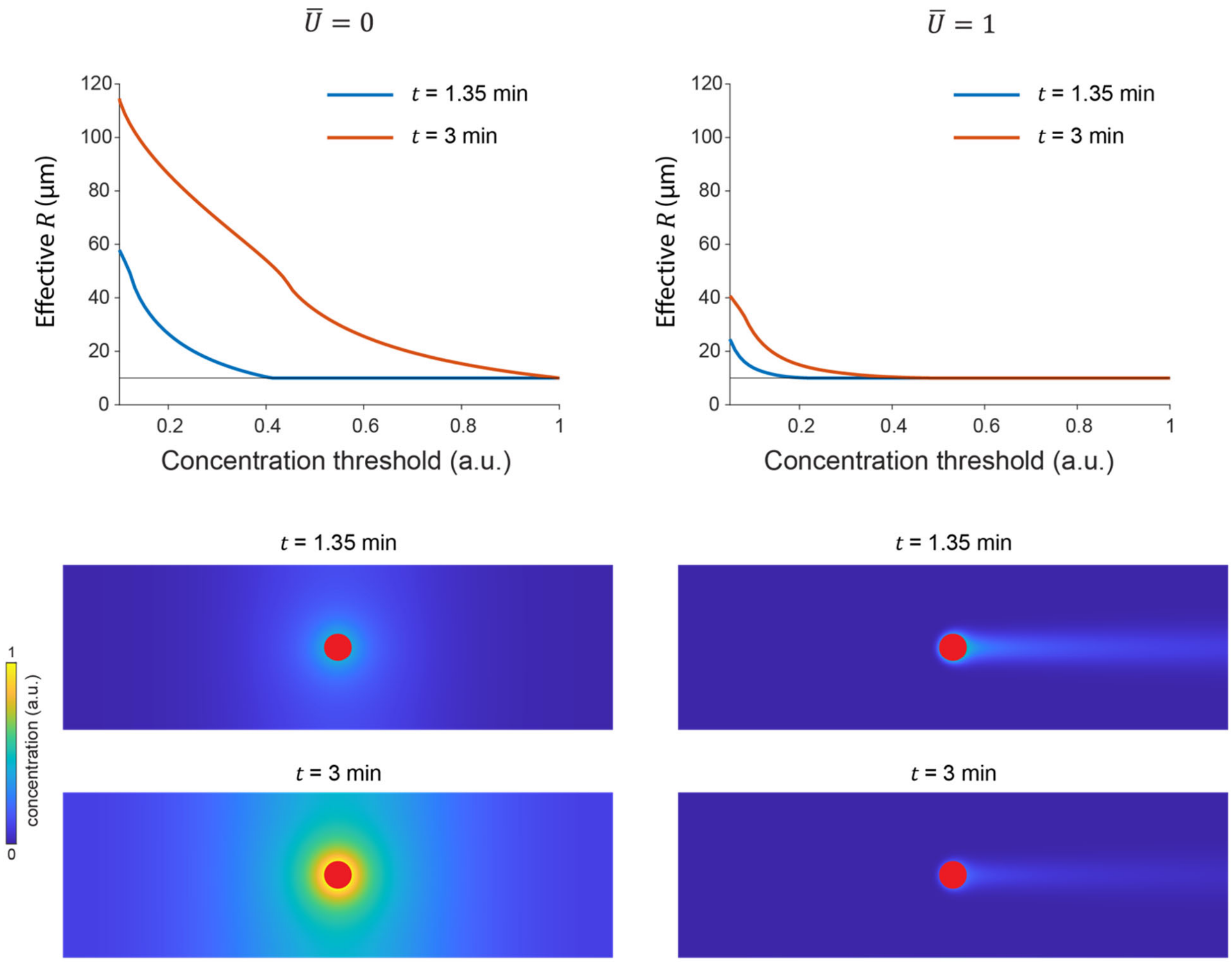


Fig. S12 | Adhesion protein concentration profiles and effective $R$ as a function of concentration threshold for protection for diffusive ($\bar{U} = 0$) and advective ($\bar{U} = 1$) cases at different physical times in simulation. All concentration profiles are depth-averaged and normalized by the value at the edge of the producer cluster at $\bar{U} = 0$ and $t = 3$ min. Each red disk corresponds to a producer cluster of radius $r_0$. The simulation shows that although $R$ increases with time in both cases, $R$ is always smaller in the advective case than in the diffusive case, and that the difference between the two cases increases with time. This strengthens our conclusion that advection reduces $R$, irrespective of the limitation on physical time scale accessible in simulation.

### Supplementary Table

**Table S1. *V. cholerae* strains used in this study**

| Strain | Genotype | Source |
|---|---|---|
| JN132 | *vpvC*$^{W240R}$ Δ*vc1807*::*P*$_{tac}$-*mScarlet-I*-Spec$^{R}$ | This study |
| JY458 | *vpvC*$^{W240R}$ Δ*bap1*Δ*rbmC* Δ*vc1807*::*P*$_{tac}$-*mNeonGreen*-Spec$^{R}$ | This study |
| JY451 | *vpvC*$^{W240R}$ Δ*vc1807*::*P*$_{tac}$-*mNeonGreen*-Spec$^{R}$ | This study |
| JN144 | *vpvC*$^{W240R}$ Δ*bap1*Δ*rbmC* Δ*vc1807*::*P*$_{tac}$-*mScarlet-I*-Spec$^{R}$ | This study |
| JY488 | *vpvC*$^{W240R}$ *bap1*-3×FLAG Δ*vc1807*::*P*$_{tac}$-*mNeonGreen*-Spec$^{R}$ | This study |
| JY489 | *vpvC*$^{W240R}$ *rbmC*-3×FLAG Δ*vc1807*::*P*$_{tac}$-*mNeonGreen*-Spec$^{R}$ | This study |
| JY459 | *vpvC*$^{W240R}$ Δ*bap1*Δ*rbmC* Δ*vc1807*::*P*$_{tac}$-*SCFP3A*-Spec$^{R}$ | This study |
| ZJ033 | *vpvC*$^{W240R}$ Δ*rbmC* Δ*vc1807*::*P*$_{tac}$-*mNeonGreen*-Spec$^{R}$ | This study |
| ZJ053 | *vpvC*$^{W240R}$ Δ*bap1* Δ*vc1807*::*P*$_{tac}$-*mNeonGreen*-Spec$^{R}$ | This study |
| JN131 | Δ*vc1807*::*P*$_{tac}$-*mScarlet-I*-Spec$^{R}$ | This study |
| JY567 | Δ*bap1*Δ*rbmC* Δ*vc1807*::*P*$_{tac}$-*mNeonGreen*-Spec$^{R}$ | This study |

**Table S2. Parameters used in the two-part competition model**

| Parameters | Symbol | Values |
|---|---|---|
| Producer growth rate | $r_{\mathrm{p}}$ | 0.75 $h^{-1}$ |
| Cheater growth rate | $r_{\mathrm{c}}$ | 1.07 $h^{-1}$ |
| Producer carrying capacity | $N_{\mathrm{p}}$ | 18.3 cells/$\mu m^2$ |
| Cheater carrying capacity | $N_{\mathrm{c}}$ | 18.3 cells/$\mu m^2$ |
| Exploitation radius | $R$ | 63 μm |
